# Dynamics of motor direction representation in the primate premotor and primary motor cortices during sensorimotor learning

**DOI:** 10.1101/2023.09.13.556461

**Authors:** Teppei Ebina, Akitaka Sasagawa, Dokyeong Hong, Rieko Setsuie, Keitaro Obara, Yoshito Masamizu, Masashi Kondo, Shin-Ichiro Terada, Katsuya Ozawa, Masato Uemura, Masafumi Takaji, Akiya Watakabe, Kenta Kobayashi, Kenichi Ohki, Tetsuo Yamamori, Masanori Murayama, Masanori Matsuzaki

## Abstract

Sensorimotor learning requires reorganization of neuronal activity in the premotor cortex (PM) and primary motor cortex (M1). However, how PM- and M1-specific reorganization occurs in primates remains unclear. We conducted calcium imaging of these areas in common marmosets while they learned a two-target reaching (pull/push) task. Throughout learning, the dorsorostral PM (PMdr) showed peak activity earlier than the dorsocaudal PM (PMdc) and M1. PMdr showed decreased representation of newly introduced (push) movement, whereas PMdc and M1 maintained high representation. Many task-related neurons in PMdc and M1 exhibited a strong preference to either movement direction. PMdc neurons dynamically switched their preferred direction, whereas M1 neurons stably retained their preferred direction. Differences in preferred direction between adjacent neurons in PMdc increased during learning. These results suggest that in primate sensorimotor learning, dynamic motor representation in PMdc converts the cognitive sensorimotor signals of PMdr to stable and specific motor representation of M1.

## Introduction

Sensorimotor transformation is a fundamental function in the brain. In our daily lives, we execute different movements according to different sensory cues. We learn sensorimotor associations in many situations and then come to stably execute the appropriate movement in each situation. In particular, for both humans and non-human primates, the execution of a variety of forelimb movements is very important, regardless of the requirement for hand dexterity. The forelimb motor cortical area is included in the dorsal part of the premotor cortex (PMd) and primary motor cortex (M1) (Nudo et al., 1996; Strick et al., 2021). In primates, the rostral part of the PMd (PMdr) connects densely with the prefrontal cortex (PFC) and is associated with cognitive processes, while the caudal part of PMd (PMdc) strongly connects with M1 and possesses corticospinal neurons that imply that it is more strongly related to motor processes than is PMdr (Abe and Hanakawa, 2009; Barbas and Pandya, 1987; Luppino et al., 2003; Picard and Strick, 2001; Wise et al., 1997). In forelimb movement, PMdr neurons show an earlier response to action-related sensory signals than do PMdc and M1 neurons, and PMdc activity follows PMdr activity after the action to be taken is determined (Cisek and Kalaska, 2005; Grafton et al., 1998). PFC and PMdr are thought to be important in the association of sensory cues with movement in sensorimotor learning (Deiber et al., 2014; Hikosaka et al., 2002; Shadmehr and Holcomb, 1997). PM and M1 neurons show sensorimotor learning-related and motor skill learning-related changes in activity (Brasted and Wise, 2004; Germain and Lamarre, 1993; Li et al., 2001; Mitz et al., 1991; Paz and Vaadia, 2004; Zach et al., 2008). However, the activities of neurons in PMdr and PMdc, or in PMdc and M1, are frequently collected as the same population when recording motor cortical activity. Thus, it is unclear how the PMdr, PMdc, and M1 motor representations differ during sensorimotor learning.

PMdc and M1 have a hallmark property in that individual neurons possess a preferred direction for reaching movements and neighboring neurons show a similar preferred direction (Amirikian and Georgopoulos, 2003; Ben-Shaul et al., 2003; Georgopoulos et al., 2007). Although this preferred direction is stable across days after motor learning (Bollimunta et al., 2021; Chestek et al., 2007; Ebina et al., 2018), it is unclear how PMdc and M1 neurons show directional preference when a new-direction movement is introduced into a reaching task and learned over dozens of days.

One-photon and two-photon calcium imaging have clarified the reorganization of M2 (which is assumed to correspond to PM) and M1 during learning of forelimb movement tasks in rodents (Makino et al., 2017; Masamizu et al., 2014). However, the rodent M2, which includes the rostral forelimb area (RFA), is responsible for a wide range of information processing, including action planning, decision making, working memory, behavioral adaptation, and grasping (Barthas and Kwan, 2017; Ebbesen and Brecht, 2017; Svoboda and Li, 2018). These functions are not spatially separated within M2, and hierarchical areas that correspond to PMdr and PMdc have not been found within M2. Thus, to understand the functions of differentiated premotor cortical areas, it is necessary to examine primate species.

Recently, calcium imaging of the motor cortex was established in common marmoset and macaque while they performed forelimb movement tasks (Bollimunta et al., 2021; Ebina et al., 2018; Kondo et al., 2018; Trautmann et al., 2021). This research revealed that the preferred direction of forelimb reaching in the superficial layer neurons in PMdc and M1 is relatively stable across days (Bollimunta et al., 2021; Ebina et al., 2018). However, it remains unclear whether the stability and spatial distribution of the preferred direction differs between PMdc and M1. The neuroanatomical structures (including PMdr and PMdc) of the marmoset brain share a high degree of similarity with other primates (Bakola et al., 2015; Burman et al., 2008; Krubitzer and Kaas, 1990), but the marmoset cerebral cortex is much smaller than that of the macaque, and has a flat and smooth surface that is suitable for calcium imaging (Kalaska, 2019; Matsuzaki and Ebina, 2020; Walker et al., 2017). In the current study, we performed wide-field one-photon calcium imaging of PMdr, PMdc, M1, sensory cortex, and posterior parietal cortex, and two-photon calcium imaging of PMdc and M1, to detect neuronal activity over a learning period of more than 2 weeks for a simple two-target reaching (pull/push) task in marmosets that were already experts in a one-target reaching (pull) task (Ebina et al., 2018). Using this imaging data, we reveal how the pull-related and push-related activity in PMdr, PMdc, and M1 changed. We also clarify how the preference direction (PD) (between pull and push) of PMdc and M1 neurons, and the PD spatial distribution, change during the learning period.

## Results

### Learning of the two-target reaching task

We trained six head-fixed marmosets (marmosets 1–6) to perform a visual-cue-triggered one-target reaching (OTR) task for 31–85 days, and then trained them to perform a two-target reaching (TTR) task over 15–43 sessions (Ebina et al., 2018) (Fig. 1A, B, and Fig. S1A, B). In the TTR training sessions, we conducted one-photon calcium imaging of marmosets 1–3 and two-photon calcium imaging of marmosets 4–6 (Fig. S1A and Table S1). In the OTR task, the animal needed to pull a pole that could move two-dimensionally in the horizontal plane with their left forelimb to move a cursor from a fixation area to a target area. This cursor appeared below the fixation area on the monitor in front of the animal (Ebina et al., 2018). The two-dimensional pole movement was directly linked to the two-dimensional cursor movement. After the cursor was moved to the target area and held within it, a sucrose water reward was delivered. In the TTR task, the animal needed to pull or push the pole to move the cursor to a target area that appeared either below or above the fixation area, respectively (Fig. 1B). As the training sessions progressed, the pull movement in the OTR task and the push movement in the TTR task were made more difficult by shortening the width of the target area, lengthening the time of cursor holding within the target area, and narrowing the permissible range of the opposite initial movement (Fig. 1C). Over the training sessions, the cursor trajectory in the rewarded trials became straight to fulfill the final threshold for the reward acquisition (Fig. 1C). We defined the rewarded trials that did not show any initial opposite movement as successful trials (see Methods for details). During the TTR task sessions, the rate of successful trials to total pull trials remained high (∼0.8), while the rate of successful push trials to total push trials increased to ∼0.8 (Fig. 1D, E). In all of the following analyses of the neuronal activity, we use only data from successful trials in those sessions with at least 10 successful pull and push trials. From these sessions, and for each animal, two or three sessions were selected as early sessions and three sessions as late sessions (Table S1). In five out of six animals, the variability of the trial-to-trial cursor trajectory compared with the averaged trajectory in the successful push trials was smaller in the late sessions than in the early sessions (Fig. 1F, G). The reaction time (the duration from the target onset to the time at which the cursor exited the fixation area) in the successful pull trials increased from early to late sessions (Fig. 1H, I).

**Figure 1.**
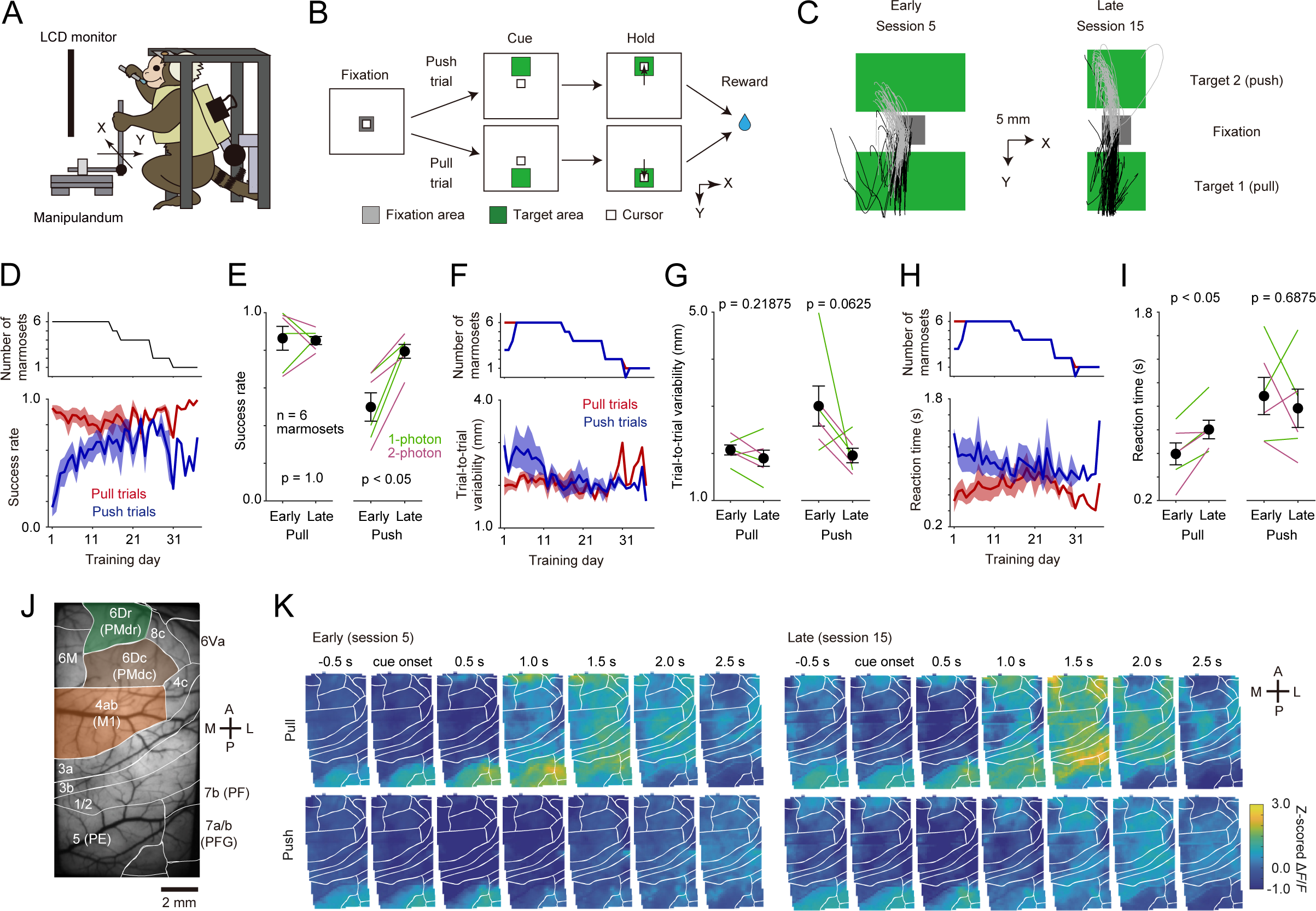
One-photon calcium imaging of the frontoparietal cortex during learning of a two-target reaching task. (A) Scheme of the task apparatus and head-fixed marmoset. (B) Two-target reaching task. After the cursor was fixed in a fixation square, a target (green) appeared randomly above or below it in each trial. When the animals pulled or pushed the pole to move the cursor downward or upward, respectively, to the target, and then held it within the target, they obtained the reward. (C) Reaching trajectories in sessions 5 and 15 of the TTR task in marmoset 1. Black and gray lines represent the trajectories during –500 to 500 ms from the movement onset for all trials with targets below (target 1; 117 trials in session 5 and 67 trials in session 15) or above (target 2; 109 trials in session 5 and 68 trials in sessions 15) the fixation square, respectively. The green rectangle and gray square represent the target rectangles and fixation square, respectively. The width of the target square gradually shortened through the sessions, and the distance between the fixation square and target 2 increased. (D) Time courses of the rates of successful trials to the total pull (red) and push (blue) trials (n = 6 animals). Shading indicates ± SEM. The number of animals on each day is shown in the top chart. (E) The rates of successful pull and push trials in the early and late sessions. Green indicates marmosets 1–3 for one-photon imaging, and purple indicates marmosets 4–6 for two-photon imaging. Black dots indicate means and bars indicate SEM. (F) Time course of the trial-to-trial variability of the pole trajectory in successful pull and push trials (n = 6). (G) The trial-to-trial variability of the pole trajectory in successful pull and push trials in the early and late sessions. (H) Time course of the reaction times in successful pull and push trials (n = 6). (I) Reaction times of successful pull and push trials in the early and late sessions. (J) The one-photon imaging window of marmoset 1. White lines indicate boundaries of cortical areas inferred from the ICMS motor map and the marmoset brain atlas. The corresponding area name and/or numbers are also shown. (K) Trial-averaged pseudo-colored z-scored Δ*F*/*F* images at 0.5 s time points between 0.5 s before and 2.5 s after the cue onset for pull and push trials in sessions 5 and 15 in marmoset 1.

### One-photon calcium imaging of wide cortical areas including PMdr, PMdc, M1, and parietal areas during learning of the TTR task

For each animal, the locations of motor cortical and other areas were inferred from intracortical microstimulation (ICMS) mapping results (Fig. S2A, B) and a stereotaxic brain atlas (Paxinos et al., 2012). During the OTR task training, adeno-associated viruses (AAVs) carrying tetracycline-inducible tandem GCaMP6s were injected (Chen et al., 2013; Hashimoto et al.; Sadakane et al., 2015) (Fig. S1A). We used GCaMP6s to obtain a fluorescence change with a higher signal-to-noise ratio than that from GCaMP6m and GCaMP6f, although the decay time constant of GCaMP6s is much slower than that of GCaMP6m or GCaMP6f (Chen et al., 2013). Thus, we focused on only the early phase of the task-related fluorescence change, which was relatively less affected by the slow decay kinetics. For one-photon imaging, the AAVs were injected into multiple sites over right cortical areas including PMdr, PMdc, M1, the somatosensory cortex, and the posterior parietal cortex (PPC) in marmosets 1–3. Then, a large cranial window (15 × 8 mm) was placed to cover these areas (Fig 1J and Fig. S2A). For two-photon calcium imaging, the AAVs were injected into PMdc and M1 in marmosets 4–6 and a rectangular cranial window (9 × 5 mm) was placed (Fig. S2B). During the imaging, the marmoset chair and the one-photon microscope were slightly tilted to introduce the excitation light into the frontoparietal cortex perpendicular to the glass window (Ebina et al., 2018). After completing one-photon imaging experiments in marmosets 1 and 2, we confirmed that muscimol injection into the right PM, M1, or PPC, but not the primary somatosensory area (S1), inhibited the pull and push movements (Fig. S2C, D). Saline injection into the right PM or M1 did not affect the task performance (Fig. S2D). During one-photon and two-photon imaging, we also recorded body movements (forelimb, hindlimb, and orofacial movements) with two cameras and extracted the movements of several body parts with the DeepLabCut toolbox (Ebina et al., 2019; Mathis et al., 2018) (Fig. S3A–D).

One-photon imaging clearly showed the dynamics of the neuronal activity (relative fluorescence change, Δ*F*/*F*) in the TTR task over wide areas (Fig. 1K). In the following analyses, the image size was down-sampled to 64 × 64 pixels to reduce the noise affecting each pixel and the calculation time. The calculation was based on the Δ*F*/*F* trace of each pixel, and the calculated values were averaged within each area of the inferred PMdr, PMdc, and M1. After the calcium imaging experiment during the TTR task sessions in marmoset 3, we confirmed that Δ*F*/*F* was not contaminated by a change in the intrinsic fluorescence reflecting hemodynamic activity (Pisauro et al., 2013) by comparing calcium-dependent fluorescence obtained with light illumination at a wavelength of 470 nm with calcium-independent fluorescence obtained with a 405 nm light (Allen et al., 2017) (Fig. S4A–D).

### The order of the timing of the peak activity of PMdr, PMdc, and M1 was stable during learning of the TTR task

Although a variety of activity patterns were detected across the PMdr to the PPC (Fig. 1K), we focused on the three motor cortical areas (PMdr, PMdc, and M1) because our aim was to reveal the spatiotemporal dynamics of PM and M1 activity during the TTR task. The activities in these three areas increased after cue onset and reached a peak around 2 s after cue onset (Fig. 2A). A transient response to the target presentation was not apparent, and most activity appeared to be movement related (Fig. 2A, B). First, we estimated the peak timing of the trial-averaged activity (*T*_peak_) for pull and push trials in the early and late sessions in marmosets 1–3 (Fig. 2C). In the *T*_peak_ map, PMdr showed the earliest peak timing in all twelve conditions (early and late sessions in successful pull and push trials in three animals) (Fig. 2D and Fig. S5A). In eleven conditions, *T*_peak_ was earlier in PMdc than in M1. When we calculated the correlation between *T*_peak_ in each area and the reaction time across the training sessions, the *T*_peak_ of PMdr and reaction time correlated in all six conditions (successful pull and push trials in three animals; Fig. 2E, F), although *T*_peak_ was slower than the reaction time. These results suggest that PMdr activity before the peak timing might be strongly related to the reaction time of the pull and push movements throughout learning, and that the peak activity flowed from PMdr to PMdc and PMdc to M1.

**Figure 2.**
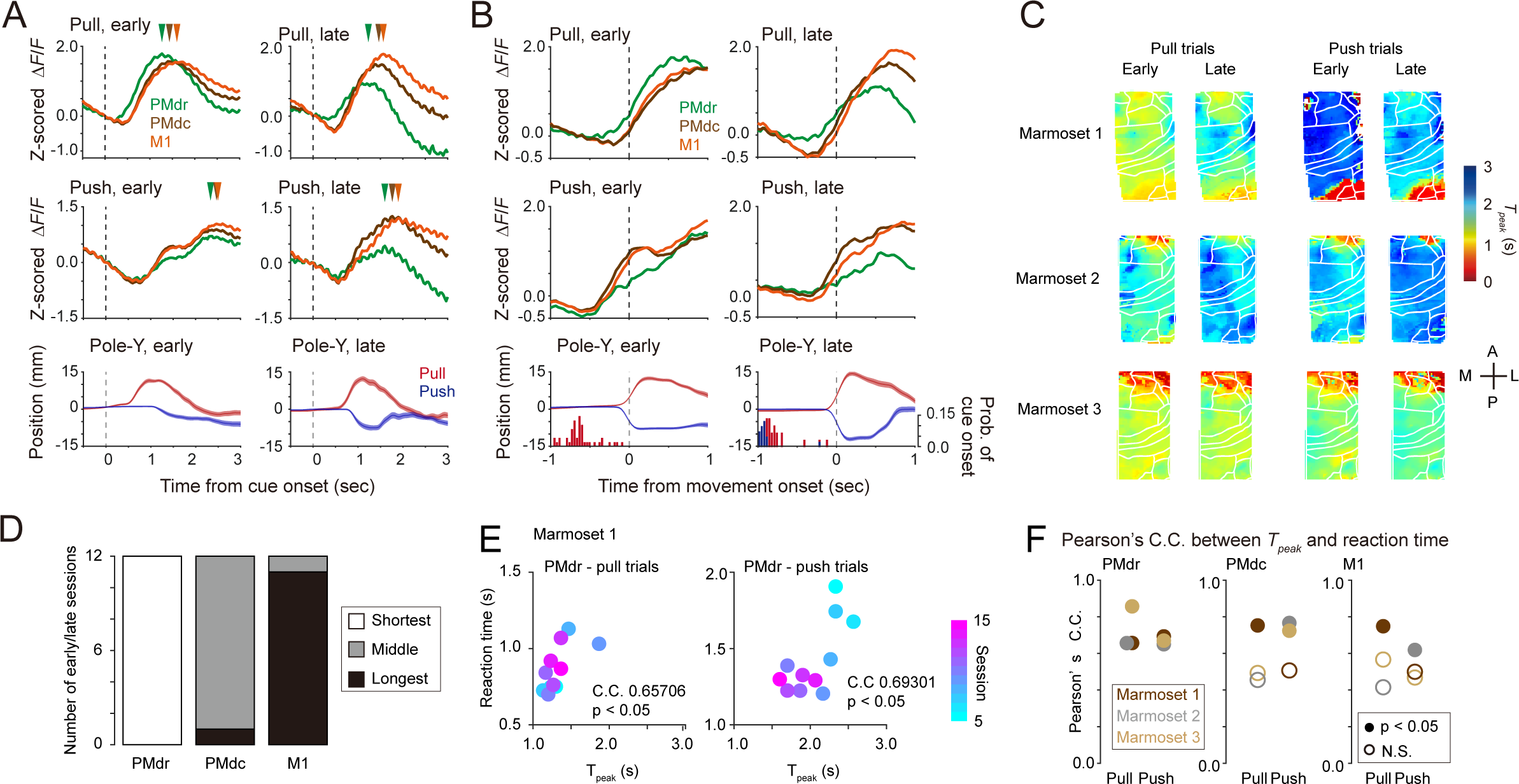
Activity changes in PMdr, PMdc, and M1 during sensorimotor learning. (A, B) Δ*F*/*F* traces of area-averaged PMdr (green), PMdc (brown), and M1 (orange) in pull (top) and push (middle) trials in early (left) and late (right) sessions of marmoset 1. In (A), the activity is aligned to the onset of the target presentation. Colored arrowheads indicate *T_peak_* of the corresponding colored traces. The bottom image shows the averaged Y-axial pole trajectory for pull (red) and push (blue) trials. In (B), the activity is aligned to the onset of the pole movement. A histogram of the cue onset timing is overlaid. (C) Pseudo-colored maps of the timing of the trial-averaged peak activity (*T_peak_*) for pull and push trials in early and late sessions of marmosets 1–3. The Tpeak at each pixel was averaged over the early or late sessions. White lines indicate the boundaries of the putative cortical areas. (D) The number of sessions with the shortest, middle, and longest *T_peak_* out of the 12 sessions for each area. (E) Plot of reaction time against *T_peak_* of PMdr for pull and push trials in all the 11 analyzed imaging sessions from marmoset 1. (F) Pearson’ s correlation coefficients between the *T_peak_* of the three areas and the reaction time for pull and push trials in marmosets 1–3 during the training period (brown, marmoset 1; gray, marmoset 2; orange, marmoset 3). Closed circles indicate that the correlation was statistically significant (p < 0.05) and open circles indicate that it was not statistically significant. The number of imaging sessions was 11, 17, and 12 for marmosets 1, 2, and 3, respectively.

### PMdc and M1 retained stronger motor representation than PMdr

Next, we examined how information on the forelimb movement was possessed by each area throughout learning. In push trials, the push movement was frequently followed by a pull movement (Fig. S3C). Since the slow decay time constant of GCaMP6s substantially hindered detection of activity changes associated with fast switches in movement direction, we focused on the initial movement after the cue presentation. To estimate the motor information of PMdr, PMdc, and M1, we decoded the cursor movement from 1 s before to 0.13 s after the movement onset that included only pull or push movement (Fig. 3A). The cursor position at each time point was predicted from the neuronal activity from 0.3 s before to 0.3 s after the time point. For each pixel of the imaging field, we calculated the cross-validated coefficient of determination (cv*R*²) to represent the prediction accuracy (Kondo and Matsuzaki, 2021; Terada et al., 2022) (Fig. 3A). Throughout learning, the area-averaged cv*R*² was much higher in PMdc and M1 than in PMdr in both early and late sessions and both types of trials, although the cv*R*² values varied across the three animals (Fig. 3B and Fig. S5B). In eleven out of the twelve conditions, cv*R*² was higher in M1 than in PMdc (Fig. 3C and Fig. S5B). PMdr showed decreased prediction accuracy for push trials, although this was not statistically significant in marmoset 2 (Fig. 3D). PMdc and M1 did not show a consistent trend in the change of cv*R*² for pull or push movements across training sessions (Fig. 3D).

**Figure 3.**
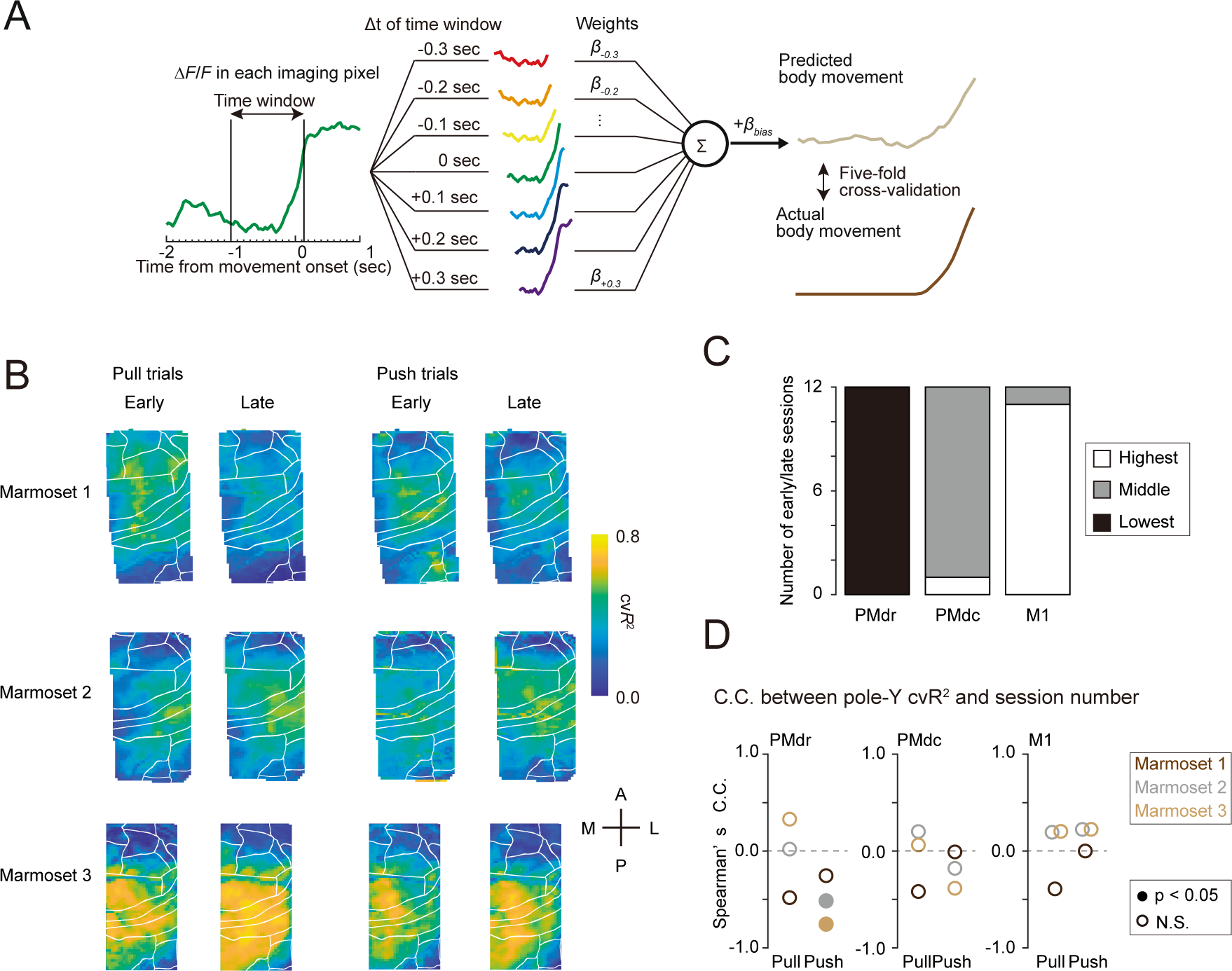
Changes in the prediction accuracy of initial movements in PMdr, PMdc, and M1 during the sensorimotor learning. (A) Decoding model using linear regression. Pole position at a given time t (from 1 s before to 0.133 s after the movement onset) was predicted from the neuronal activity data every 0.1 s for 0.3 s before and after each time point. (B) Pseudo-color map of prediction accuracy (cv*R*^2^) for pull and push trials in early and late sessions in marmosets 1–3. The prediction accuracy was averaged over the early or late sessions. (C) The number of sessions with the highest, middle, and lowest cv*R*^2^ out of the 12 sessions for each area. (D) Spearman’ s rank correlation coefficient between cv*R*^2^ and imaging sessions for marmosets 1–3. Closed circles indicate that the correlation is statistically significant (p < 0.05) and open circles indicate that it is not statistically significant. The number of imaging sessions was 11, 17, and 12 for marmosets 1, 2, and 3, respectively.

The activity of the imaged PM and M1 might reflect task-related orofacial movements rather than the contralateral forelimb movement, although the orofacial movement should be more represented in the ventral PM and ventral M1 (Burish et al., 2008). However, when the pupil diameter, Z-axial eye movement, and lick rate were predicted from the neuronal activity in each area, the cv*R*² was smaller than the cv*R*² of the initial pole movement in most cases, and the prediction accuracy of any of the three orofacial movements did not consistently increase or decrease (Fig. S5C–H). The Y-axial pole movement showed constant strong correlations with the Y-axial left hand and Y-axial left elbow movements, but did not strongly correlate with the Z-axial eye movement, pupil diameter, lick rate, right hand, right elbow, or bilateral knees throughout the training sessions (Fig. S5I). Thus, the orofacial, left forelimb, and hindlimb movements did not have a large effect on the measured activity in PMdr, PMdc, and M1. These results suggest that PMdc and M1 retained the representation of both pull and push forelimb movements, while the representation of the newly introduced push movement in PMdr was weak and decreased throughout learning. In the following analyses and experiments, we focus on the PMdc and M1, which appeared to carry stronger motor information than PMdr.

### The change in preferred direction was larger in PMdc than in M1

We next examined the change in the PD in PMdc and M1 pixels. We defined the preferred direction index (PDI) as (cv*R*² in pull trials – cv*R*² in push trials) / (cv*R*² in pull trials + cv*R*² in push trials). A PDI of 1 or –1 indicates that only a pull or push movement was represented, respectively. A PDI of 0 indicates that pull and push movements were represented equally. There was no apparent millimeter-scale segregation between the high PDI sub-areas and low PDI sub-areas that were common across the three animals in either the early or late sessions (Fig. 4A). More than 80% of pixels in PMdc and M1 showed a PDI within the range of –0.2 to 0.2 in both early and late sessions, and these pixels showed cv*R*^2^ of > 0.3 for both pull and push movements (Fig. 4B). This indicates that many sub-areas strongly showed both pull- and push-related activity in both early and late sessions. However, the degree of the change in PDI in each pixel from the early to late sessions differed between PMdc and M1. Compared with M1, PMdc showed larger changes in PDI from early to late sessions, while the sign of the area-averaged PDI in PMdc was not consistent in early or late sessions across the three marmosets (Fig. 4A, C). The change in PDI in each pixel (ΔPDI_EL_) in M1 from early to late sessions was distributed around zero in all three animals, whereas ΔPDI_EL_ in PMdc deviated from zero, although the direction of the deviation was not consistent across the animals (Fig. 4D). These results suggest that more sub-areas altered their PD in PMdc than in M1. Thus, the manner of the motor representation during the training sessions differed between PMdc and M1.

**Figure 4.**
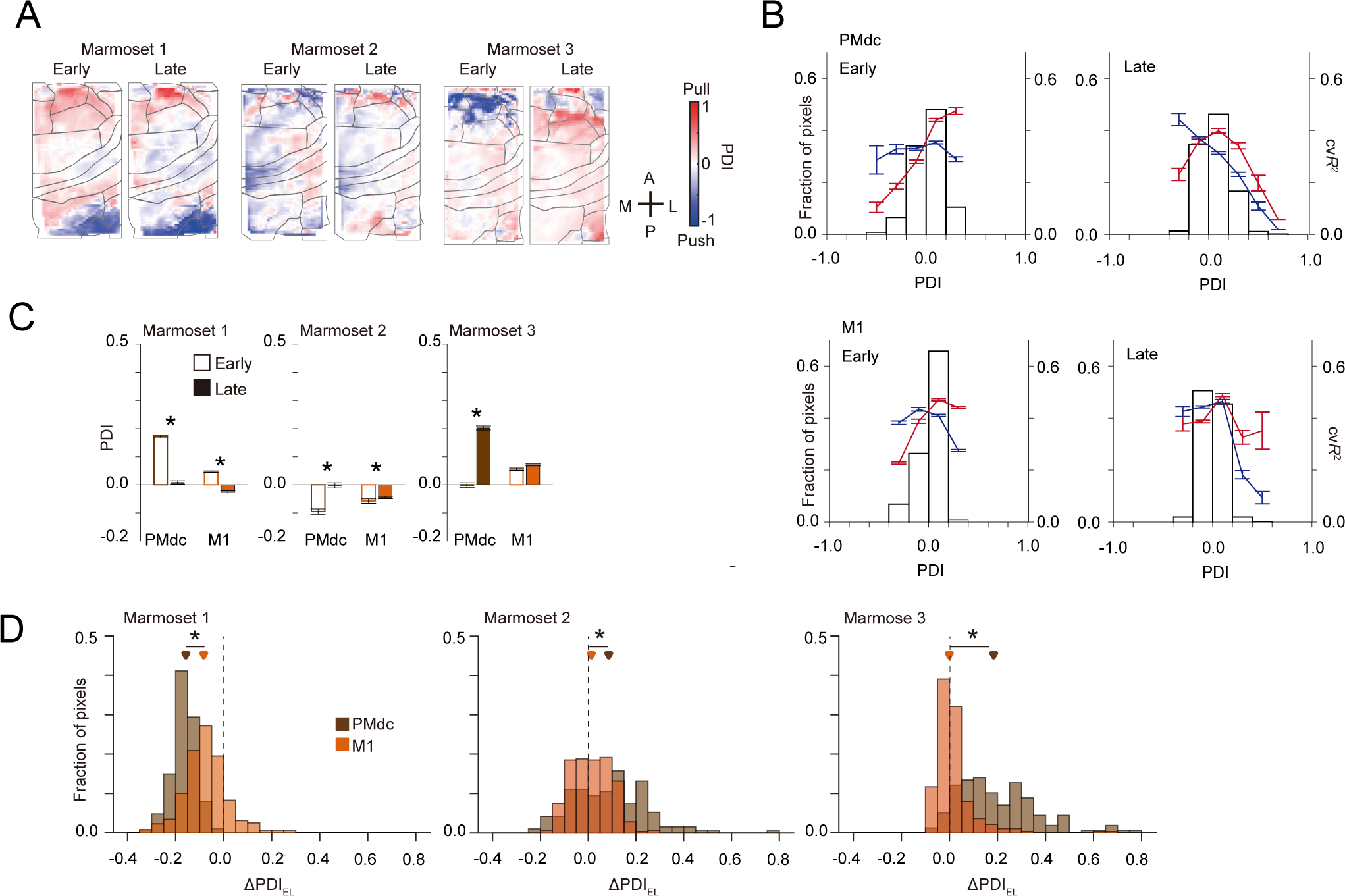
Changes in preferred direction of PMdc and M1. (A) Pseudo-color maps of PDI in early and late sessions for marmosets 1–3. The pixels with cv*R*^2^ of < 0.02 in both pull and push trials were not analyzed. (B) Histogram of the PDI (bars) of pixels in early (left) and late (right) sessions in PMdc (top) and M1 (bottom). The cv*R*^2^ averaged within each bin for pull (red) and push (blue) trials is overlaid. (C) Area-averaged PDI of PMdc and M1 in early and late sessions for marmosets 1–3. *p < 0.05, Wilcoxon sign-rank test. (D) Histograms of ΔPDI_EL_ of PMdc (brown) and M1 (red) in marmosets 1–3. The brown and red triangles indicate the median of the PDI distribution of PMdc or M1, respectively. *p < 0.05, Wilcoxon rank-sum test.

### Wide field-of-view two-photon calcium imaging after learning revealed that M1 neighboring neurons showed a more similar preferred direction than PMdc neighboring neurons

The sub-areas that represented both pull and push movements might reflect the fact that individual neurons showed similar activity between pull and push trials, or that neurons with different PDs were intermingled within the sub-areas. To examine the PD distribution of individual neurons over PMdc and M1 after learning, we applied wide field-of-view two-photon microscopy with a 3 × 3 mm field of view (FOV) (which was first developed for mouse brain imaging (Ota et al., 2021)) to a marmoset that was trained to perform the TTR task (marmoset 7; Fig. S6A, B). We expanded the space between the large objective lens and the sample stage, and to make the optical axis perpendicular to the cranial window, we set the head-fixed marmoset sitting in the chair so that it was substantially tilted (Fig. S6C).

The FOV was so large that parts of PMdc and M1 could be simultaneously imaged (Fig. 5A). We used a constrained non-negative matrix factorization (CNMF) (Pnevmatikakis et al., 2016) to extract active ROIs with denoised Δ*F*/*F* as active neurons. We detected 15 961 active neurons from six imaging sessions (in 4 days) in marmoset 7 (10 510 PMdc and 5451 M1 neurons; Fig. 5B– D). The session-averaged rates of the successful pull and push trials were 0.781 ± 0.047 and 0.640 ± 0.03 (n = 4 days), respectively. The variability values of the trial-to-trial pole trajectory in the successful pull and push trials were 1.36 ± 0.06 mm and 1.65 ± 0.04 mm, respectively. The reaction times in the successful pull and push trials were 709.4 ± 35.5 ms and 917.9 ± 56.4 ms, respectively. As for the pixels of the one-photon imaging data, the cv*R*^2^ for pull and push movements and the PDI were calculated for each active neuron (Fig. 5E). When neurons had a cv*R*^2^ of > 0.02, they were defined as pull-related or push-related neurons (Fig. 5F and Fig. S6D). Their proportions ranged from 0.15–0.25 and were slightly higher in M1 than in PMdc (Fig. 5G). In both pull and push trials, the cv*R*^2^ of the motor-related neurons was higher in M1 than in PMdc (Fig. 5H). The peak timing of the activity of movement-related neurons was earlier in PMdc than in M1 (Fig. 5I–K). This result is consistent with the results of the population activity obtained with one-photon imaging (Fig. 2D).

**Figure 5.**
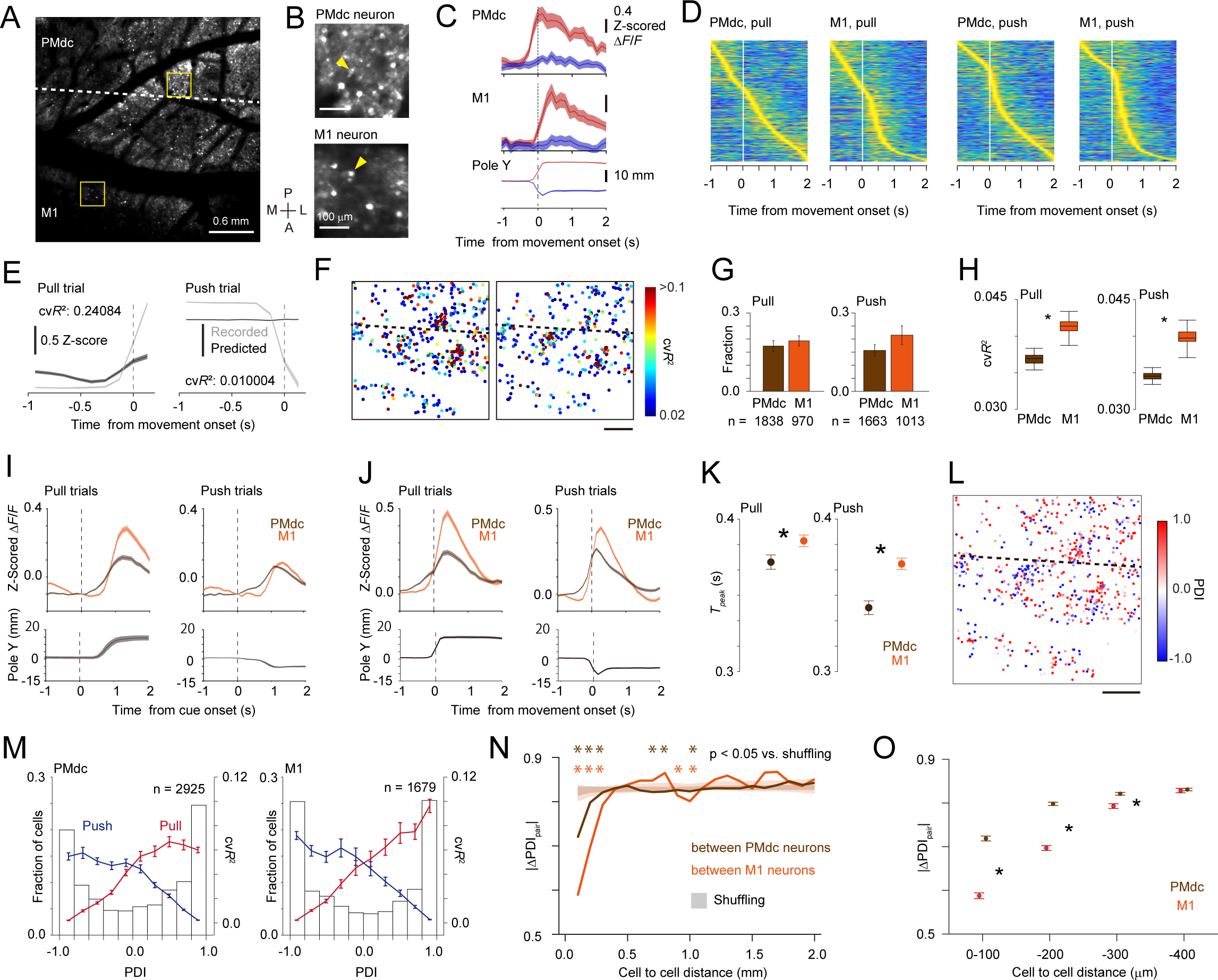
Large-field two-photon calcium imaging of PMdc and M1 after learning. (A) A representative frame-averaged two-photon image of PMdc and M1. The dotted line indicates the putative border between PMdc and M1. Scale bar, 600 μm. (B) Expanded images of the boxed regions in (A). (C) Trial-averaged movement-onset-aligned activity of the arrowed PMdc (top) and M1 (middle) neurons in (B) for pull (red) and push (blue) trials. The bottom plot shows the trial-averaged movement-onset-aligned Y-axial pole trajectory. (D) Activities of all active neurons in PMdc and M1 that were aligned to the onset of the pole movement for pull and push trials (n = 10510 PMdc neurons and 5451 M1 neurons). The neurons were ordered according to the timing of the maximum activity during –1 to +2 s from the movement onset. (E) The trial-averaged pole trajectory (gray) and the pole trajectory predicted from the activity of the PM neuron shown in (B). (F) Pseudo-color map of the cv*R*^2^ of pull/push-related neurons in the FOV shown in (A). The cv*R*^2^ of each neuron for pull (left) and push (right) trials is pseudo-colored. (G) Proportions of PMdc and M1 neurons with cv*R*^2^ of > 0.02 in pull and push trials. n = 6 sessions. The total numbers of pull-related neurons were 1838 in PMdc and 970 in M1. The total numbers of push-related neurons were 1663 in PMdc and 1013 in M1. (H) The cv*R*^2^ of PMdc and M1 neurons with cv*R*^2^ of > 0.02 for pull and push trials. Box plots represent the 95% confidence intervals of the median cv*R*^2^ of each group. The confidence interval was calculated by a bootstrap with 1000 repetitions. (I) Time course of the averaged activity of pull-related and push-related neurons in PMdc and M1 aligned to the onset of the target presentation. Trial-averaged activity is averaged over neurons. Bottom, the averaged trace of the Y-axial pole trajectory. (J) Time course of the averaged activity of pull-related and push-related neurons in PMdc and M1 aligned to the onset of the movement. Trial-averaged activity is averaged over neurons. Bottom, the averaged trace of the Y-axial pole trajectory. (K) *T_peak_* of pull-related and push-related neurons in PM and M1 for the pull and push trials, respectively (n = 1838 in PM and 970 in M1 for pull trials, n = 1663 in PM and 1013 in M1 for push trials). (L) Pseudo-color map of PDI in the FOV shown in (A). (M) Histogram of the PDI (bars) of neurons in PMdc and M1. The cv*R*^2^ averaged within each bin for pull (red) and push (blue) trials is overlaid. (N) |ΔPDIpair| for pairs of PM neurons (brown) or pairs of M1 neurons (red) against the cellular distance. Shading indicates 95% of shuffled data. Asterisks indicate that the original data exceeded 95% of the values of the corresponding shuffled data (p < 0.05). (O) |ΔPDIpair| for a cellular distance of < 400 μm. *p < 0.05, Wilcoxon rank-sum test.

Next, we examined the PDI of each neuron. Within the 3 × 3 mm FOV, there was no apparent segregation in the millimeter-scale areas of pull and push movements, but there were small scattered clusters for each direction (Fig. 5L). In both PMdc and M1, many neurons showed a high PD; ∼40% of neurons in PMdc and M1 showed PDI of < –0.8 or > 0.8 (Fig. 5M). The proportion of neurons with a PDI of –0.2 to 0.2 was only ∼10% in both areas (Fig. 5M). As PDI increased, cv*R*^2^ for the pull movement increased and cv*R*^2^ for the push movement decreased in both areas. Thus, individual neurons with strong movement-related activity in PMdc and M1 showed clear directional preference. The distributions of PDI and cv*R*^2^ of individual neurons differed substantially from the distributions of PDI of individual pixels that were imaged by one-photon imaging (Fig. 4B). These results suggest that many neurons showed high directional preference and that neurons with different PDs are intermingled in PMdc and M1.

When we calculated the absolute value of the difference in PDI between a pair of neurons (|ΔPDI_pair_|), the |ΔPDI_pair_| at a cellular distance of < 300 μm was significantly smaller than the |ΔPDI_pair_| when the cellular distance was shuffled in both M1 and PMdc (Fig. 5N). |ΔPDI_pair_| at a cellular distance of < 300 μm was significantly smaller in M1 than in PMdc (Fig. 5O). Thus, in M1, the neurons with similar PD tended to cluster more strongly in a local region of a few hundred micrometers than they did in PMdc.

### Chronic two-photon calcium imaging during learning revealed that PMdc neurons changed PD more dynamically than did M1 neurons

From the results so far, we assumed that any FOV of ∼0.6 × 0.6 mm in PMdc and M1 included both pull-related and push-related neurons to some extent, and that some of these neurons might change their activity during the TTR training sessions. Then, while marmosets 4–6 sat in the slightly tilted chair learning the TTR task, we conducted two-photon imaging of PMdc and M1 neurons with a microscope whose scanning head could be tilted and whose FOV was ∼0.6 × 0.6 mm (Ebina et al., 2018). We analyzed three FOVs in PMdc L2/3 and three FOVs in M1 L2/3 for all three animals (Fig. S2B and Fig. S3D). We pooled the data for each area and were able to pursue 398 PMdc neurons and 336 M1 neurons in at least one session of each of the early and late sessions (Fig. 6A, B). Among these pursued neurons, the neurons that showed cv*R*^2^ of > 0.02 (Fig. S7A, B) for pull or push movement were defined as movement-related neurons (pull-related neurons or push-related neurons, respectively). When we compared cv*R*^2^ between early and late sessions for each pursued neuron (Fig. 6C), there were neurons in both PMdc and M1 that showed increased cv*R*^2^, and those that showed decreased cv*R^2^*. We defined the increase or decrease neurons as neurons that showed an increased or decreased cv*R*^2^ of more than 0.02 from the early to late sessions, respectively. The proportion of decrease neurons was larger than that of increase neurons in both areas (Fig. 6D). When cv*R*^2^ was averaged over all pursued neurons except for those that were unrelated to either movement in both early and late sessions (black in Fig. 6C), the cv*R*^2^ for push movement in all three FOVs in PMdc decreased from the early to late sessions, although the decrease was not significantly different in one FOV (Fig. 6E). By contrast, the change directions in the cv*R*^2^ for pull movement in PMdc, or in the cv*R*^2^ for pull and push movements in M1, were not consistent across FOVs (Fig. 6E, F). When the population activity of these neurons was used to predict the initial pole movement along the Y axis, the direction of the change in the prediction accuracy for either movement type from the early to late sessions was not consistent across the three animals (Fig. 6G, H). This is consistent with the result obtained from one-photon imaging (Fig. 3D), and suggests that while the populations of PMdc and M1 neurons did not consistently change their prediction accuracy for either pull or push movement during the training sessions, a subset of PMdc neurons did show decreased prediction accuracy for the push movement.

**Figure 6.**
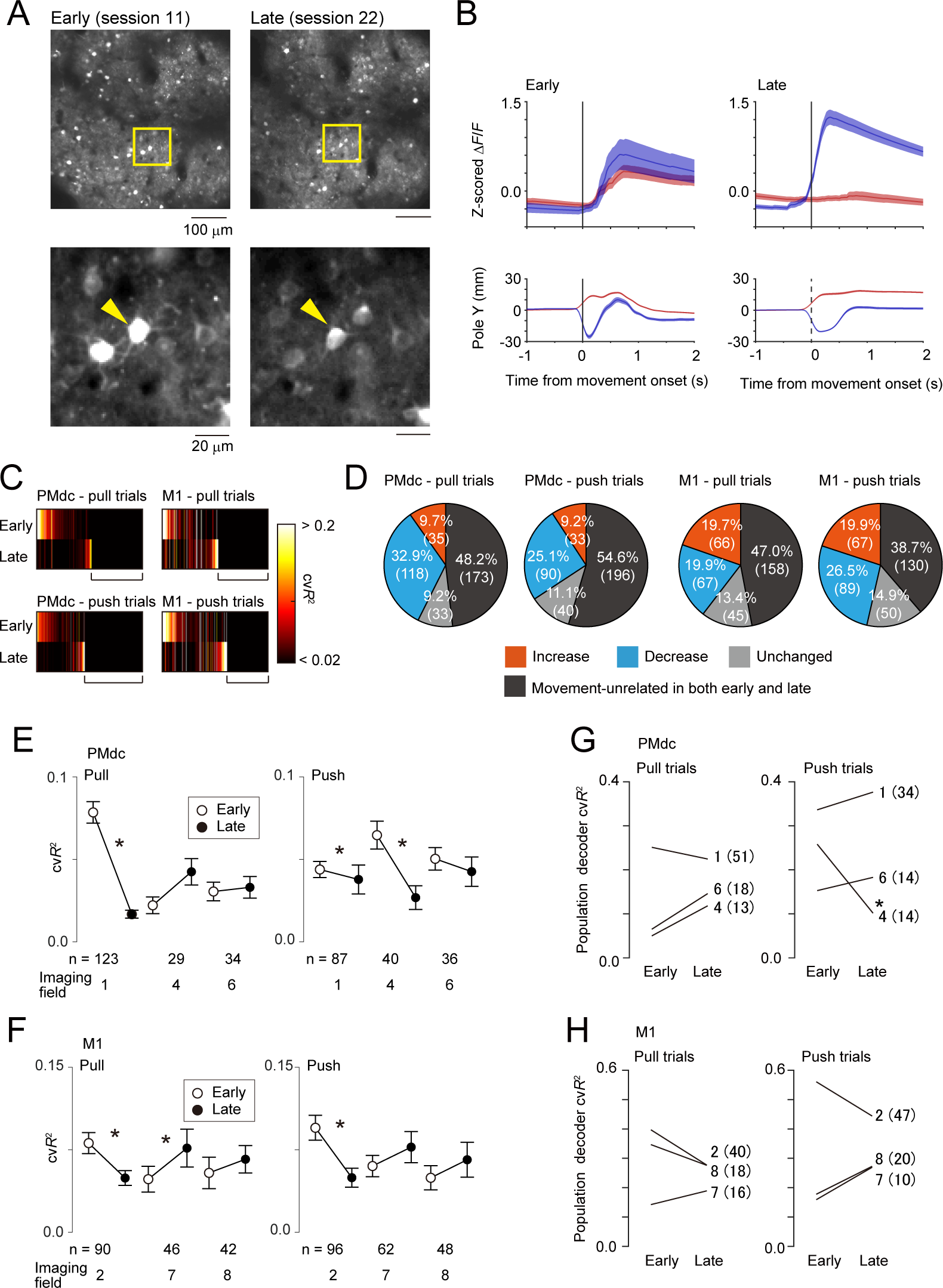
Changes in the prediction accuracy of the initial pole movement in individual PMdc and M1 neurons during the sensorimotor learning. (A) Top, representative images of the same FOV in sessions 11 (early) and 22 (late) of marmoset 4. Bottom, expanded images of the boxed regions shown in the top image. (B) Trial-averaged activity of the arrowed neurons in (A) for pull (red) and push (blue) trials. Bottom, averaged traces of Y-axial pole trajectory. (C) The cv*R*^2^ of all pursued PM and M1 neurons for pull and push trials in the early and late sessions (n = 359 pursued neurons in PMdc and 336 pursued neurons in M1). Black horizontal bars indicate the pursued neurons that were not related to the pull or push movement in either early or late sessions (movement-unrelated). (D) Proportions of the increase, decrease, unchanged, and movement-unrelated neurons in PMdc and M1 for pull and push trials. Numbers in parentheses represent the numbers of neurons. (E, F) The cv*R*^2^ of the pursued neurons for pull and push trials in each imaging field of PMdc (E) and M1 (F) in early and late sessions. *p < 0.05, Wilcoxon sign-rank test. For pull trials, the cv*R*^2^ of pursued neurons that were pull-related in the early sessions and the cv*R*^2^ of those that were pull-related in the late sessions were used, while for push trials, the cv*R*^2^ of pursued neurons that were push-related in the early sessions and the cv*R*^2^ of those that were push-related in the late sessions were used. The numbers of neurons that were used and the imaging field # (Fig. 2SB and Table S1) are also shown. (G, H) The cv*R*^2^ of the population decoders in PMdc (G) and M1 (H). Each number indicates the imaging field #. The numbers in parentheses represent the numbers of neurons used for the decoder. The number of neurons used for the decoder was the minimum number of neurons across the early and late sessions for each imaging field. *p < 0.05, the 97.5 percentile of the differences in cv*R*^2^

Finally, we examined whether the PD of individual neurons changed more in PMdc than in M1, as predicted from the results of the one-photon imaging. The distributions of PDI in the early and late sessions were similar; the proportion of neurons with a PDI of < –0.8 or > 0.8 was high in both PMd and M1, although the PDI distribution was biased to either positive or negative values within each FOV (Fig. 7A, B and Fig. S8). By contrast, the distribution of the difference in PDI between the early and late sessions (ΔPDI_EL_) was broader in PMdc neurons than in M1 neurons (Fig. 7A, B), and |ΔPDI_EL_| was larger in PMdc neurons than in M1 neurons (Fig. 7C). These results suggest that even at the single-neuron level, the directional preference changed more dynamically in PMdc than in M1 during learning. In addition, |ΔPDI_pair_| at any cellular distance of < 300 μm in PMdc differed between the early and late sessions; |ΔPDI_pair_| was ∼0.6 in the early sessions and ∼0.8 in the late sessions (Fig. 7D). This suggests that the heterogeneity of PD within the PMdc local circuit increased over the training sessions. By contrast, in M1, |ΔPDI_pair_| did not differ between the early and late sessions (Fig. 7E). Consistent with this, the wide field-of-view two-photon microscopy results obtained from marmoset 7, |ΔPDI_pair_| at a cellular distance of < 100 μm in the late sessions was smaller in M1 than in PMdc (PMdc, 0.82 ± 0.03, n = 325 pairs, M1, 0.67 ± 0.02, n = 739 pairs, p < 0.01, Wilcoxon rank-sum test). These results suggest that in comparison with M1 neurons, PMdc neurons changed PD more dynamically at both single-neuron and local-network levels.

**Figure 7.**
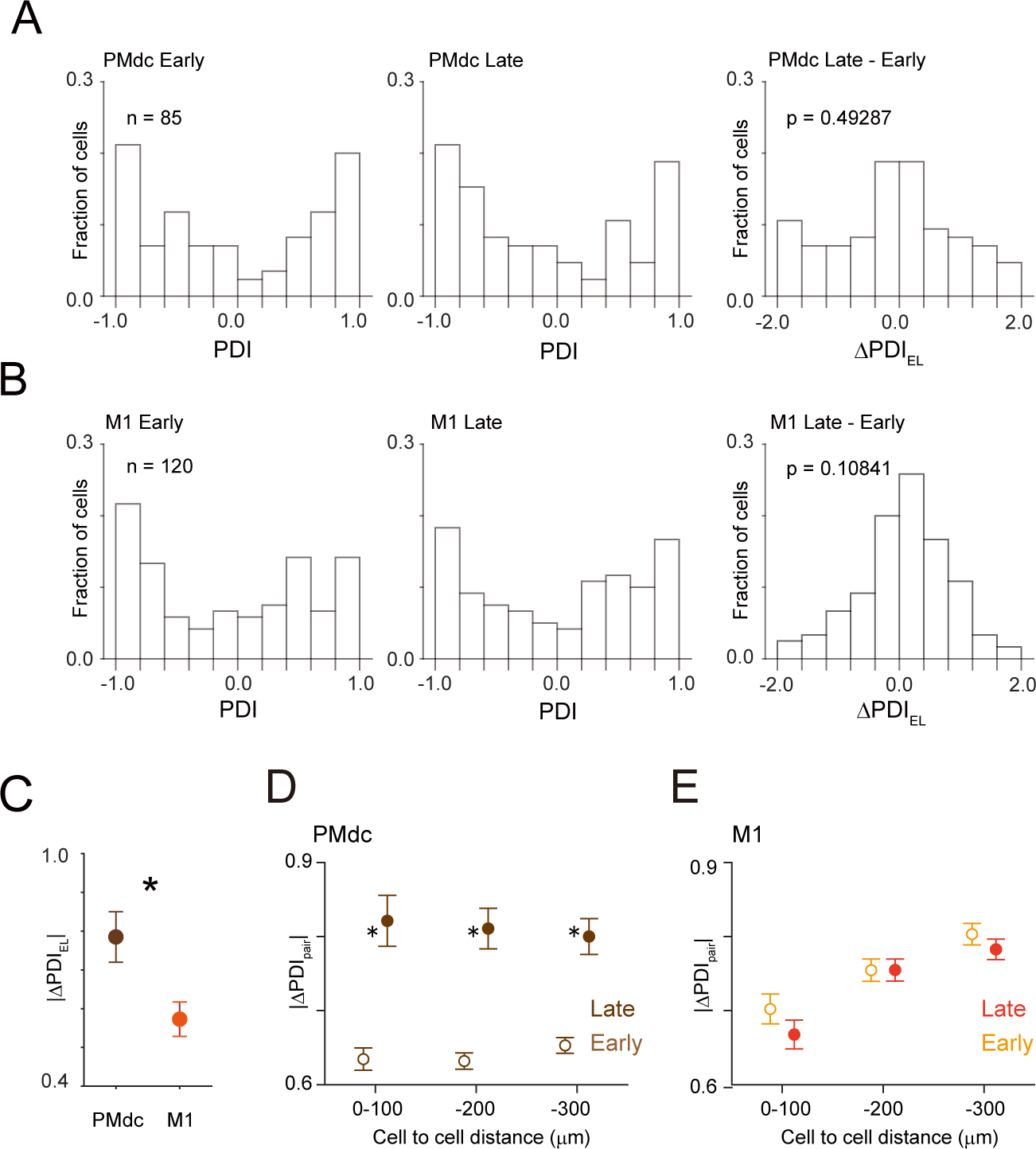
Changes in the PD of individual PMdc and M1 neurons during the sensorimotor learning. (A, B) Histograms of the PDI of the pursued neurons in the early (left) and late (middle) sessions and ΔPDI_EL_ (right) in PMdc (A) and M1 (B) neurons. Only pursued neurons that were pull-related and/or push-related in both early and late sessions were included in this analysis. (C) |ΔPDI_EL_| of the PMdc (brown, n = 85) or M1 (red, n = 120) neurons shown in A or B, respectively. *p < 0.05. Wilcoxon rank-sum test. (D, E) |ΔPDI_pair_| for pairs of pursued neurons in PMdc (D) and M1 (E) against the cellular distance in the early (open symbols) and late (closed symbols) sessions. Neurons that were pull-related and/or push-related in the early and late sessions were used in the calculation of |ΔPDI_pair_| for the early and late sessions, respectively. *p < 0.05, Wilcoxon rank-sum test comparing the |ΔPDI_pair_| between the early and late sessions.

## Discussion

In the present study, we applied calcium imaging to task-performing marmosets to reveal the reorganization of the motor cortex during the sensorimotor learning process. In particular, one-photon calcium imaging demonstrated the distinct activity of PMdr and PMdc, and two-photon calcium imaging demonstrated the differences in the change and distribution of PD between PMdc and M1 L2/3 neurons during learning.

Wide-field one-photon imaging revealed earlier peak activity in PMdr than in PMdc, while the motor representation was stronger in PMdc than in PMdr. We also detected a decrease in the push representation during learning in PMdr, but not in PMdc. The newly introduced target above the fixation should require attention because it did not appear in the OTR task. To associate the newly introduced target with the push movement in the TTR task, PMdr might need to have push-related activity in the early sessions, activity that might then attenuate as the learning progressed. Both PMdr and PMdc should be related to the sensorimotor transformation, and PMdc should also be related to the motor execution. The latter motor-related component in PMdc activity might be unrelated to the reaction time, so that the reaction time for both pull and push movement during training sessions might be more correlated with the timing of the peak activity in PMdr than with that in PMdc. The current task, in which the animal did not need to retain the sensory memory before decision making and movement onset, is different to the tasks in many previous studies in which an instructed delay period was set (Cisek and Kalaska, 2005; Kurata and Wise, 1988).

Nevertheless, the current results are consistent with many previous studies using electrophysiological recording in macaque: PMdr is strongly associated with cognitive processes based on visual information, whereas PMdc is closely related to motor processes, and PMdr activity precedes PMdc activity (Cisek and Kalaska, 2005; di Pellegrino and Wise, 1991; Sayegh et al., 2013). The PMdr-PMdc order of the activity reflects the fact that the caudal part of the PPC is associated with visual processing of movement and projects more strongly to PMdr than to PMdc (Andersen and Buneo, 2002; Burman et al., 2014), although we did not analyze the activity in the PPC.

Wide-field one-photon imaging demonstrated that the pull and push motor representation of the area-averaged PMdc and M1 did not change substantially during the training sessions. Two-photon calcium imaging demonstrated that a subset of L2/3 neurons in PMdc and M1 increased their motor representation while other neurons decreased theirs, with the prediction accuracy reflected by the ensemble activity showing no substantial change. These results are consistent with those of our previous study in mice. In that study, there were a subset of neurons that increase their motor representation and other neurons that decrease theirs in the forelimb M1 L2/3 during learning of the forelimb lever-pull task, but the prediction accuracy reflected by the population activity did not substantially change (Masamizu et al., 2014). Thus, in the superficial layer, a subset of neurons that flexibly change the amount of motor information they carry may contribute to motor learning in both primates and rodents.

However, the PM and M1 of macaques and humans frequently show expansion of the learned-movement-related area and increase in the learned-movement-related activity (Karni et al., 1995; Mitz et al., 1991; Nudo et al., 1996; Sanes and Donoghue, 2000). Although the reaching movement to the target should be novel for the marmosets, grasping the pole by the left hand and applying force (pull or push) to the pole should be familiar to them because their home cages had fences that could be grasped, and therefore a subset of neurons might have already formed a high preference for the push direction. Since we only estimated the prediction accuracy of the initial phase of the movement, we might not have detected neuronal activity reflecting the fine improvement in the push movement that was newly introduced in the TTR task. We previously reported context-dependent reorganization with fine movement proficiency in mouse M1 L2/3 (Terada et al., 2022). Another possibility was that large changes in push-related activity might have occurred in the earliest stage in which the number of successful trials was less than 10, a stage that we did not analyze. In macaque PM and M1, activities related to learning to associate new sensory cues to predetermined movements rapidly emerge within a session (Mitz et al., 1991; Paz and Vaadia, 2004; Zach et al., 2008). Alternatively, substantial reorganization might occur in the deep layer. When electrical recording in macaque shows activity change during learning, the recorded neurons generally originate from the deep layer. In mice, the activity dynamics differ between M1 L2/3 and L5 neurons during learning of the lever-pull task (Masamizu et al., 2014). Endoscopy imaging and three-photon imaging could potentially be used to measure the activity of deep layer neurons in the marmoset (Bollimunta et al., 2021; Kondo et al., 2018; Ouzounov et al., 2017).

Wide-field one-photon imaging showed no apparent segregated millimeter-scale area in PMdc and M1 for pull and push movements. In optogenetic stimulation of marmoset M1, the pull and push directions of the forelimb movement were separately mapped along the medial-lateral axis (Ebina et al., 2019), mapping that should reflect the activity of deep layer neurons. The superficial layers might integrate more different types of motor information than the deep layer, as discussed above. By contrast, wide field-of-view two-photon imaging revealed that many movement-related neurons possessed a strong preference for the reaching direction, and the similarity of PD between pairs of neurons was statistically significant at a cellular distance of < 300 μm in M1 and PMdc. In macaque, neighboring motor cortex neurons (at a cellular distance of < 400 μm; Smith and Fetz, 2009) show strong synaptic linkages (Lee et al., 1998; Smith and Fetz, 2009; Stark et al., 2008). Thus, the clustering of neurons that are related to the same movement direction is a fundamental self-organizing property in non-human primates (Amirikian and Georgopoulos, 2003; Asanuma and Rosén, 1972; Ben-Shaul et al., 2003; Stark et al., 2008). However, we have not imaged the same area at different depths, nor the activity related to other movement directions. Three-dimensional mapping of reaching in eight directions is necessary to clarify whether the local clusters that we detected are parts of functional columns, and how much local size is represented in all directions (Amirikian and Georgopoulos, 2003; Georgopoulos et al., 2007).

Across both early and late sessions, the PD of M1 neurons was more stable than that of PMdc neurons. Although we examined activity for only pull and push directions, our results do not contradict the previous finding that macaque M1 neurons did not substantially change their preferred direction within a session of a visuomotor association task with eight-way reaching (Paz et al., 2003). After learning, the PD similarity between neighboring neurons was higher in M1 than in PMdc. It did not change from the early to late sessions in M1 neurons, whereas it decreased in PMdc. Thus, the PD became heterogeneous in PMdc, even within the local area of a few hundred micrometers. The reaction time for pull movement increased during learning (Fig. 1I), which might be related to learning of the specific association between the cue and pull movement that was not necessary for the OTR task with only one reaching direction. Because of the association between the push target and push movement and the re-association between the pull target and pull movement, the PDs of PMdc neurons might be bidirectionally changed. These results suggest a principle of spatiotemporal patterns of sensorimotor transformation as follows: PMdc neurons within individual sub-areas flexibly process different sensorimotor associations and their interactions, convert the sensory signal to the motor initiation signal, and transfer the signal to clusters of M1 neurons that have been specifically differentiated to accurately execute individual movements.

PMdr and PMdc were substantially different in the manner of their motor representation, as were PMdc and M1. Thus, it is apparent that the motor cortex is more differentiated in the marmoset than in the rodent (Barthas and Kwan, 2017; Chen et al., 2017). This difference is probably related to differences in the repertoires of forelimb movements between primates and rodents. Primate ancestors evolved the ability to move the forelimb independently of other body parts, and to differently move left and right forelimbs for grasping and leaping in arboreal life (Nyakatura, 2019). In addition, primates should need to flexibly associate many sensory targets, such as fruits and insects, with a variety of behavioral repertoires (Schiel et al., 2010). This requirement would lead to differentiation of the M2 (more specifically, the RFA) seen in rodents to PMdr and PMdc (and also the pre-supplementary and supplementary motor cortices) in primates. The application of wide-field one-photon and wide field-of-view two-photon imaging methods to the neocortex of behaving marmosets offers promise for further understanding the cortical mechanisms of the complex sensorimotor transformations occurring in primates.

## Limitations

In the present study, we did not image neuronal activity during learning of the OTR task. Thus, we were not able to detect the dynamics related to the emergence of the association between the target cue and forelimb movement, nor the process for improving the pole-pull movement. It is not clear how the learning order from the pull to push movements affected the neuronal dynamics during the imaging sessions. The marmoset brain is lissencephalic, and it is therefore difficult to determine the borders of the PMdr, PMdc, and M1 according to landmarks on the cortical surface such as the arcuate sulcus. Although we conducted ICMS in individual animals to account for the effects of individual differences, this was not sufficient to accurately identify the borders. Registration of the imaging field to a brain atlas with MRI would be useful to more accurately identify the areas (Ose et al., 2022).

## Supplemental information

The supplemental information includes eight figures and one table.

## Author contributions

T.E. and M.Ma. designed the experiments. T.E., A.S., R.S., Ke.O., and Ka.O. conducted the experiments. T.E., A.S., D.H., and M.K. analyzed the data. T.E., Y.M., and S.T. optimized the task apparatus for the experiments. Y.M., M.U., M.T., A.W., K.K., K.O., and T.Y. designed and prepared AAVs. M.Mu. developed the wide field-of-view two-photon microscope system. T.E., A.S., and M.Ma. wrote the paper, with comments from all authors.

## Acknowledgments

We thank Y. Hirayama, M. Hirokawa, Y. Takahashi, and A. Yamamoto for animal handling. We thank T. Ode for helping to build the wide field-of-view two-photon microscope system. This work was supported by Grants-in-Aid for Scientific Research on Innovative Areas (17H06309 to M.Ma., 19H05307 to T.E., and 21H00302 to T.E.), for Transformative Research Areas (A) (22H05160 to M.Ma. and 23H04977 to T.E.), for Scientific Research (A) (19H01037 and 23H00388 to M.Ma.), for Scientific Research (B) (20H03546 to T.E.), and for Young Scientists (A) (17H04982 to Y.M.) from the Ministry of Education, Culture, Sports, Science, and Technology, Japan; AMED (JP23dm0207069 to M.Ma.; JP23dm0207001 to T.Y., M.Mu., and M.Ma.; JP18dm0207027 to M.Ma.; JP23dm0107150 to M.Ma.; JP23dm0207085 to T.E. and M.Ma.); and the Tokyo Society of Medical Sciences (to T.E.). This work was also supported by the program for Brain Mapping by Integrated Neurotechnologies for Disease Studies (Brain/MINDS) from AMED under Grant number JP21dm0207111.

## Materials & Correspondence

Correspondence and requests for materials should be addressed to Masanori Matsuzaki.

## Methods

### Animals

All experiments were approved by the Animal Experimental Committee of the University of Tokyo and the Animal Care and Use Committees of the RIKEN Center for Brain Science. Seven laboratory-bred common marmosets (*Callithrix jacchus*) were used in the present study. Their age ranged from 20–68 months and their weight from approximately 250–350 g when the habituation started. Marmosets 1–3 were used for one-photon imaging throughout motor learning, marmosets 4– 6 were used for two-photon imaging throughout the learning, and marmoset 7 was used for wide field-of-view two-photon imaging after more than 140 sessions of learning. Marmosets 1 and 6 were females and the others were males. All seven marmosets were kept under a 12:12-hour light-dark cycle. None of them were used for any other experiments prior to the present study.

### Virus production

The AAV plasmid of human synapsin I promoter (hSyn)-tetracycline-controlled transactivator 2 (tTA2) was constructed by subcloning the DNA fragments containing hSyn and tTA2 into pAAV-MCS (Agilent Technologies, CA, USA). The AAV vector was produced as described previously (Ebina et al., 2018, 2019; Konno and Hirai, 2020). For the generation of pAAV-TRE-GCaMP6s-P2A-GCaMP6s-WPRE (tandem GCaMP6s), gene fragments of GCaMP6s and P2A were obtained from pGP-CMV-GCaMP6s-WPRE (Addgene: #40753) and pAAV-hSyn1-GCaMP6s-P2A-nls-dTomato (Addgene: #51084), respectively (Hashimoto et al.). pAAV-TRE-GCaMP6f-WPRE was used as a template for this plasmid. AAV plasmids were packaged into AAV serotype 9 using the AAV Helper-Free system (Agilent Technologies). In brief, pAAV vector, pRC9, and pHelper plasmids were transfected into HEK293 cells. Seventy-two hours after transfection, AAV2/9 particles were purified using the AAV Purification kit (Takara, Siga, Japan). The AAV solution was concentrated to the optimal volume by centrifugation using an Amicon Ultra-4 100k centrifugal filter unit (Millipore). The number of genomic copies was quantified with intercalating dyes (Thermo Fisher Scientific, MA, USA) and two sets of primers for WPRE or hGHpA genes using LightCycler 480 (Roche, Basel, Switzerland). The final titration of the AAV was estimated as relative quantitation according to a calibration curve calculated from the known numbers of copies of AAV plasmids.

### Surgical procedures

All the surgeries and viral vector injections were carried out under aseptic conditions described previously (Ebina et al., 2018, 2019). Each marmoset was placed in a stereotaxic instrument (SR-6C-HT; Narishige, Tokyo, Japan); 0.8–4.0% isoflurane anesthesia at a flow rate of 1 L/minute was maintained; and the saturation of percutaneous oxygen (SpO₂), pulse rate, and rectal temperature were monitored throughout surgery. In the perioperative period, the following medicines were intramuscularly (i.m.) administered: ampicillin (16.7 mg per kg of body weight) as an antibiotic, carprofen (4.4 mg/kg) as an anti-inflammatory agent, and maropitant (1000 mg/kg) as an antiemetic. In order to avoid dehydration, Ringer’s solution (10 mL) and riboflavin (vitamin B₂) were subcutaneously infused.

In the head plate implantation procedure, we depilated and sterilized the scalp, incised it with an external application of lidocaine, and removed the connective tissue to expose the skull. After six to seven small screws were anchored to the skull, a headplate was attached to the skull with universal primer (Tokuyama Dental, Tokyo, Japan), dual-cured adhesive resin cement (Bistite II or Estecem II; Tokuyama Dental), and dental resin cement (Superbond; Sun Medican, Siga, Japan). The task training under head fixation started more than a week after the head plate implantation.

### ICMS

After sufficiently long task-learning sessions (Fig. S1), craniotomy and durotomy were carried out under anesthesia and the medications described above were administered with additional intramuscular administration of dexametazone (0.5 mg/kg) and subcutaneous administration of D-mannitol (2 g/kg) to prevent cerebral edema. The exposed cortex was covered with silicone elastomer (Kwik-Sil; World Precision Instruments, FL, USA) and further covered with dental resin cement (Superbond; Sunmedical). To identify the boundary between the primary motor cortex (M1) and premotor cortex (PM), we conducted ICMS in a similar way to that described previously (Ebina et al., 2019), 2–7 days after the craniotomy for marmosets 3–7, and after the imaging experiments were finished for marmosets 1–2 (Fig. S1). During the ICMS, we anesthetized each marmoset with ketamine (initial dose 15 mg/kg; additional dose 5 mg/kg) and xylazine (initial dose 0.75 mg/kg; additional dose 0.25 mg/kg), and administered atropine (0.050 mg/kg) for sialoschesis to prevent respiratory obstruction. A silver reference electrode was immersed in the cerebrospinal fluid on the exposed cerebral surface, and then a tungsten microelectrode with an impedance of 0.5 MΩ and a diameter of 100 μm was inserted into the cerebral cortex to a depth of 1.5 or 1.8 mm. Twelve or fifteen 0.2 ms cathodal pulses of 333 Hz were applied. We increased the stimulation currents from 10 to 100 μA in steps of 10 μA until a body movement was detected.

### AAV injection

Mineral oil (Nacalai Tesque, Kyoto, Japan) was back-filled into a Hamilton syringe (25 μL) and a quartz pipette with an outer tip diameter of approximately 30 μm (Sutter Instruments, CA, USA). Then, the viral solution containing rAAV2/9-TRE3 promoter-tandem GCaMP6s (1.55 × 10¹³ or 1.36 × 10¹³ vector genomes [vg]/mL) and rAAV2/9-hSyn-tTA2 (3.9×10¹³ vg/mL) was front-loaded with a syringe pump (KDS310; KD Scientific, MA, USA). The viral solution was vertically injected into each site at a depth of 500 μm from the cortical surface at a rate of 0.10 μL/minute to a total amount of 0.50 μL. Then, the pipette was maintained in place for 5 minutes before being slowly withdrawn. After injections into multiple sites, the exposed cortical surface was covered with a rectangular glass window of 15 × 8 mm for marmosets 1–3, 9 × 5 mm for marmosets 4–6, and a circular window with a diameter of 5.5 mm for marmoset 7.

### Task behaviors

The marmosets were seated on the task apparatus while wearing a marmoset jacket (Ebina et al., 2018). The task consisted of five steps, as described in Ebina et al. (2018): primary training without head fixation, pole-pull task without head fixation, pole-pull task with head fixation, OTR task, and TTR task (Fig. S1A). In the primary training, the animals were given water and feed while seated in the chair to acclimate them. After the animals were sufficiently acclimated, we started training them in the pole-pulling task, in which a sucrose water reward was delivered when they pulled a pole with their left forelimb under a head-unfixed condition. After the animals were able to perform the pole-pull task for approximately 60 minutes, the pole-pull task with head fixation was started.

For the OTR and TTR tasks, a seven-inch liquid crystal display (LCD) monitor was placed 10–17 cm in front of the animal. Pulling and pushing of the pole moved the cursor downward and upward, respectively. Moving the pole to the right and left moved the cursor to the right and left, respectively. Each trial of the OTR task began with a holding period during which the pole was moved to the center position by a spring force of 0.25 N. That is, the cursor was moved to the fixation area. If the cursor stayed within the fixation area for 1400–2100 ms (randomly chosen for each trial), the fixation square and the spring force disappeared, and the green target rectangle appeared below the fixation area signaling the beginning of the reaching period. The target rectangle was presented during this period of 10–20 s. Animals were rewarded when they pulled the pole to move the cursor to the target within the reaching period and the cursor was held within the target rectangle for a set time (rewarded trials). In rewarded trials, the color of the target changed to white, and the white target was presented for 500 ms before it disappeared. The reaching period, holding time, distance between the fixation and target, and size of the target rectangle changed during the training sessions. When the holding time was 400 ms or longer and the rate of the rewarded trials to the total trials was 70% or higher, the animals were considered to be experts and the TTR task started. In the TTR task, either of two green target rectangles appeared: a pull target below or a push target above the fixation square. The animals needed to move the cursor to the target within the allotted time and hold it within the target for the allotted time to receive 30–100 μL of sucrose water as a reward. The final parameters were 1000–1200 ms fixation period, 100 ms holding time within the target, an upper limit of 20 mm movement in the opposite direction, 8 × 8 mm fixation area size, 16 × 16 mm target size, and 14 mm distance between the centers of the fixation and target. In the TTR task sessions, we defined the early and late sessions as follows. Among the imaging sessions with at least 10 successful pull and push trials each, the early sessions were the first two or three imaging sessions before the 11th training session or those in which the rate of successful push trials was less than 0.5 (for two early sessions in marmoset 4), and the late sessions were the last three sessions in which both the rate of successful pull and push trials was more than 0.5 (Table S1).

The trial-to-trial variability of the pole trajectory for each session was defined as the mean of the root mean square deviations (RMSDs) of the X and Y coordinates of individual trajectories from those of the trial-averaged trajectory. For each trial, the RMSD was calculated as 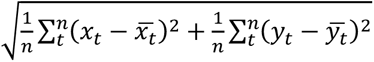, where *n* is the number of time points during the period from −500 to +200 ms of the pole movement onset, and *x*_*t*_/*y*_*t*_ and *x̅*_*t*_/*y̅*_*t*_ are the X/Y coordinates of the trajectory in the trial and trial-averaged trajectory at time point *t*, respectively. The marmosets performed the task 1–5 days per week, over which their body weight was maintained at approximately 90% of their normal level by restriction of food and water. On the off-duty days, the food and water restrictions were weakened to allow them to return to their normal weight. The task events were controlled by LabVIEW software (National Instruments, TX, USA). The task data were sampled at 1 kHz.

To monitor body movements during the task performance, two CMOS cameras (DMK33UP1300, ImagingSource, Taipei, Taiwan) were placed at 35° and 90° angles from the front of marmosets 1 and 2, with single focal length lenses with f-numbers of 35 mm and 3 mm, respectively. Images of 320 × 240 pixels were acquired at a frame rate of 100 Hz. For marmosets 3– 6, the cameras were placed at a 35° angle from the front of the marmoset with a single focal length lens (f = 3 mm) and a varifocal lens (f = 2.7–12.0 mm, Spacecom, Japan). For these marmosets, the pixel resolution and frame rate were changed to 480 × 480 pixels and 50 Hz to increase the prediction performance of the body movement tracking with DeepLabCut. For marmoset 7, the cameras were placed at 35° and 90° angles from the front of the marmoset with a single focal length lens (f = 35 mm and 3 mm, respectively). The pixel resolution and the frame rate of the images were set to 480 × 480 pixels and 30 Hz, respectively.

### One-photon imaging

One-photon imaging in marmosets 1 and 2 was conducted with a variable zoom microscope (Axio Zoom.V16; Carl Zeiss, Jena, Germany) equipped with an air objective lens (Plan-NEOFLUAR Z 2.3 ×; numerical aperture 0.5; Carl Zeiss) and a FOV of 12.6 × 12.6 mm. The marmoset chair was rotated by 0°–6° in the anterior-posterior direction and 0°–12° in the lateral-medial direction. For marmosets 1 and 2, the microscope was not tilted. For marmoset 3, the microscope was tilted by 5°– 15° in the anterior-posterior direction. These adjustments allowed the introduction of excitation light into the frontoparietal cortex perpendicular to the glass window (Ebina et al., 2018). A fluorescence light source (HXP 200 C; Carl Zeiss) and a filter set (38HE, Carl Zeiss; 470/40 nm excitation filter, 495 nm dichroic mirror, and 525/50 nm emission filter) were used for the imaging experiments. The intensity of the emitted excitation light was 5.0–6.5 mW. During the imaging, the animal’s head and the objective lens were covered with lightproof cloths to shut off possible stray light. A scientific CMOS camera (Sona; Andor Technology) with a resolution of 2048 × 2048 pixels was used as a photodetector, and the imaging was conducted at a frame rate of 30 Hz. Each series of imaging data consisted of 5400 frames (three minutes). Two-to-seventeen series of imaging data were acquired during a session. In marmoset 3, one-photon imaging was conducted with a custom-made zoom-variable microscope equipped with an air objective lens (Plan-NEOFLUAR Z 1.0 ×; numerical aperture 0.25; Carl Zeiss). The FOV size was set to 14.6 × 14.6 mm. A fluorescence light source (M470L5; Thorlabs) and a filter set were used for the imaging experiments. The excitation light intensity under the objective was 6.0 mW. A scientific CMOS camera (ORCA-Fusion; Hamamatsu Photonics) with a resolution of 2304 × 2304 pixels was used as a photodetector and the images were acquired at a frame rate of 30 Hz. Each series of imaging data consisted of 5400 or 10 800 frames (3 or 6 minutes). Two-to-seventeen series of imaging data were acquired during a session.

One-photon imaging to estimate the contamination of hemodynamic signals in the calcium imaging was also conducted in marmoset 3. An illumination light at 405 nm (M405L4, Thorlabs) was used to detect non-calcium-dependent fluorescence (Allen et al., 2017; Musall et al., 2019). The excitation light intensity under the objective was 9.0 mW for 470 nm and 3.0 mW for 405 nm. The images were acquired at a frame rate of 40 Hz. The excitation wavelength was switched from frame to frame, resulting in a 20 Hz frame rate for each excitation light.

### Two-photon imaging during motor learning

For marmosets 4–6, we conducted two-photon imaging with a custom-built two-photon microscopy system (Olympus, Tokyo, Japan) that is described in Ebina et al. (2018). A femtosecond pulse laser (Femtolite FD/J-FD-500; pulse width, 191–194 fs; repetition rate, 51 MHz; wavelength, 920 nm; IMRA, MI, USA) was introduced to the microscope scanning head through a neodymium-based fiber so that X-Y scanning was possible without any tilt effect on the microscope body. The excitation beam was then passed through a dichromic mirror (transmission wavelength range, 800– 1300 nm; reflection wavelength range, 400–755 nm) and a water immersion objective lens (XLPLN10XSVMP; numerical aperture, 0.6; working distance, 8 mm; Olympus). The intensity of the excitation beam under the objective lens was 35.0–65.0 mW. The fluorescence signal from the cortical tissue was reflected by the dichromic mirror and delivered to a cooled high-sensitivity photomultiplier tube through a liquid light guide with an infrared-cut filter (32BA750 RIF; wavelength range, 400–760 nm; Olympus).

The optical axis of the objective lens was inclined by an angle of 5.5°–15° and the chair was rotated horizontally to make the optical axis perpendicular to the cranial window. Bowl-shaped aluminum foil was attached to the head plate on the animal with silicone elastomer (Kwik Cast, World Precision Instruments; Dentsilicone-V, Shofu, Japan) and the space above the animal’s head was covered with light-shielding cloths. Two-to-ten imaging series were acquired at a frame rate of 30 Hz for 3 minutes (5400 frames) using FV30S-SW software (Olympus). The resolution of the imaging field was 512 × 512 pixels, with a pixel length corresponding to 1–1.2 μm. Body movements were recorded in the same way as for one-photon imaging.

### Wide field-of-view two-photon imaging

Imaging was conducted with a wide field-of-view microscopy system (Nikon, Tokyo, Japan) (Ota et al., 2021). This was equipped with a large objective lens (dry objective, 0.8 numerical aperture, 56 mm pupil diameter, Strehl ratio ∼0.99 over the FOV, working distance of 4.5 mm, 35 mm focal length) and large-aperture (14 mm^2^ aperture) gallium arsenide phosphide photomultipliers (GaAsP PMTs; R15248-40, Hamamatsu Photonics, Japan) with high current output (50 µA). A Ti:sapphire laser (Chameleon Vision-S Coherent Inc) tuned to 920 nm was introduced into a pre-chirper and then led to the resonant and galvanometric mirrors, and to the pupil of the objective lens through the scan tube lenses. The laser power under the objective lens was 60–90 mW. Emitted light was collected through 775 nm and 560 nm long-pass dichroic mirrors and 515–565 nm and 600–681 nm band-pass emission filters with GaAsP PMTs. A series of images were acquired at a frame rate of 7.5 Hz and a resolution of 2048 × 2048 pixels using Falcon software (Nikon). The total imaging duration was 10 minutes (4600 frames) for each imaging session.

The anterior-posterior (AP) axial-angle-adjustable stage was placed above the goniometer, and the lateral-medial-axial-angle and AP-angle adjustable marmoset chair was placed above the stage. The marmoset chair was used to restrain the body and to fixate the head of the task-behaving marmoset (Ebina et al., 2018). The marmoset chair and the stage were tilted to position the focal plane of the objective lens parallel to the glass window placed on the cortical surface of the behaving marmoset. The angle of the stage was adjusted every session. The space between the objective lens and the animal’s head was covered with aluminum foil to shield the objective from sprayed light. Before the surgery for the virus injection and glass window placement, marmoset 7 was habituated to perform the TTR task in the tilted chair in the microscopy environment.

### Pharmacological inactivation

To examine the effect of neuronal inactivation on task behavior, muscimol was injected into appropriate brain areas of marmosets 1 and 2 after all imaging experiments were finished. Before the first injection, the glass cranial window was replaced with a silicon-based window (thickness of 100 μm, 6-9085-12, AS-ONE Corporation, Japan), through which a microinjection needle was penetrated. Using a needle with an outer diameter of approximately 60 μm made from a quartz pipette and joined to a Hamilton syringe (25 μL), 0.25 μL of 5 μg/μL muscimol was injected at a rate of 0.10 μL/minute at a depth of 500 μm in one of the following cortical areas: PM, M1, the somatosensory area, and the posterior parietal area (Fig. S2C). As a control experiment, the same amount of saline was injected on other days. Two hours after the injection, the marmosets performed the TTR task that they had already learned. The interval between each injection experiment was more than or equal to 1 day. In our previous study (Terada et al., 2022), the lateral spread of muscimol in the mouse neocortex was estimated to be at most 2.5 mm. Taken together with the fact that the inactivation effects on the forelimb movement were not detected following muscimol injection into the somatosensory cortex in the current study, the muscimol spreading should be mainly limited to within the injected area. In the injection into the PM, muscimol was injected around the putative border between PMdr and PMdc (Fig. S2C), so that both PMdr and PMdc would be inactivated.

### Processing of one-photon imaging data

Tangential drifts in the imaging were removed with a finite Fourier transform algorithm (de Castro and Morandi, 1987) and the data were down-sampled from 2048 × 2048 or 2304 × 2304 pixels to 256 × 256 pixels. For each pixel of the down-sampled image, Δ*F*/*F*(*t*), the relative change in fluorescence at a time point *t*, was defined as (*F*(*t*)–*F*₀(*t*))/*F*₀(*t*), where *F*(*t*) is the fluorescence intensity at a time point *t*, and *F*₀(*t*) is the 8th percentile of *F*(*t*) across *t* ± 15 s. Data for the initial and end 15 s periods, and data that included time with missing video data, were excluded. The position of each cortical area across the experimental sessions was registered with the NoRMCorre program (available at https://github.com/flatironinstitute/NoRMCorre).

Imaging data were further down-sampled from 256 × 256 pixels to 64 × 64 pixels. For each pixel, the z-scored Δ*F*/*F* traces were calculated and normalized within each session. The z-score of each pixel was denoised by singular value decomposition (SVD), and the imaging data in the analyzed trials were concatenated in the time direction to create a data matrix. Next, SVD was performed on this matrix to decompose the pixel × time matrix data into multiple spatiotemporal components. Of these, the first 200 components explained more than 90% of the variance. Therefore, the original data were reconstructed from these 200 elements to remove the noise from the data (Musall et al., 2019). The reconstructed data (denoised Δ*F*/*F* data) were used for the following analyses.

The timing of the peak activity (*T*_peak_) was calculated from the successful trials in which the movement onset occurred more than 0.5 s and less than 4.0 s after the cue onset to exclude possible trials in which the animal quickly moved the pole by a random guess without looking at the target properly, or trials in which the animal might not have looked at the target. The z-scored trace was processed with Savitzky-Golay filtering with two orders and 15 frames (= 0.5 s). For each type of pull and push trial, the timing of the maximum peak of the trial-averaged z-scored trace within 3 s after the cue onset was defined as *T*_peak_ for each pixel. If the pixel had no peaks within 3 s after the cue onset, it was removed from the analysis.

To estimate the motor representation of each pixel, we constructed a decoding model to predict the pole movement from the neuronal activities, referring to a previous study (Tanaka et al., 2018). We applied a multiple linear regression model to the Δ*F*/*F* trace to predict the z-scored Y-axial trajectory of the pole from 1 s before to 0.13 s after the movement onset. The model was fitted separately for pull and push trials. In this process, we first down-sampled pole trajectory data at 30 Hz by averaging the positions during the acquisition of each imaging frame. We then aligned the Δ*F*/*F* trace to the movement onset, defined as when the cursor was moved outside the fixation square, and set seven time windows that were shifted by 100 ms (corresponding to three frames) in a 300 ms range before and after each time point. The predicted Y-axial trajectory at time *t*, *ŷ(t)*, was expressed using the following formula:

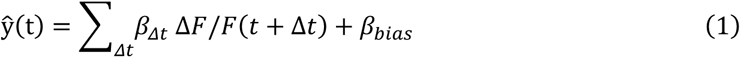

where Δ*t* was set to 0, ± 0.1, ± 0.2, or ± 0.3 s, *β*_Δ*t*_ is the coefficient at Δ*t*, and *β*_*bias*_ is the bias term. *ŷ(t)* was calculated by 5-fold cross validation. To estimate the decoder performance, the cross-validated coefficient of determination (cv*R*²) was used as a measure of prediction accuracy. We calculated the coefficient of determination as the square of the correlation coefficients between the observed and predicted Y-axial pole trajectory.

The PDI for each pixel was defined as (cv*R*² in pull trials – cv*R*² in push trials)/(cv*R*² in pull trials + cv*R*² in push trials). A PDI of 1 or –1 indicates that only pull or push movement was represented, respectively. To ensure the denominator was not too small, the pixels that showed cv*R*^2^ > 0.02 in at least one type of successful pull and push trial were used for the PDI calculation. However, all PMdc and M1 pixels in marmosets 1–3 showed cv*R*^2^ > 0.02, and therefore all these pixels were used for the PDI calculation.

When the hemodynamic-signal contamination to the fluorescence signal in calcium imaging was estimated, the Δ*F*/*F* from the data obtained with the 405 nm light (violet light-excited Δ*F*/*F*, violet-excited Δ*F*/*F*) was z-scored and smoothed using a moving average filter with a time window of 400 ms. For each pixel, the z-scored Δ*F*/*F* from the data obtained with the 470 nm light (blue light excited Δ*F*/*F,* blue-excited Δ*F*/*F*) was linearly fitted with the smoothed z-scored violet-excited Δ*F*/*F*. Then, the smoothed z-scored violet-excited Δ*F*/*F* multiplied by the weights used for fitting was subtracted from the z-scored blue-excited Δ*F*/*F* to calculate the hemodynamic corrected Δ*F*/*F* (Fig. S4). As shown in Fig. S4B–D, the blue-excited Δ*F*/*F* and the corrected Δ*F*/*F* mostly overlapped, indicating that the hemodynamic contamination in the present one-photon imaging dataset was subtle. Therefore, we did not correct the one-photon calcium imaging dataset for hemodynamic contamination.

### Processing of two-photon imaging data

Motion correction for two-photon imaging data was conducted with the same protocol used for the one-photon imaging data. Then, active neuronal somata were extracted using a CNMF algorithm (Pnevmatikakis et al., 2016). We defined the active neurons as those ROIs whose automatically extracted activities reached the following criteria: a minimum spatial component size of 50 μm^2^; a minimum spatial component ellipticity of 0.5; a minimum signal-to-noise ratio of the estimated component of 2. For each active neuron, we calculated the detrended relative fluorescent change Δ*F*/*F* = (*F*–*F*_0_)/*F*_0_, where *F*_0_ is the eight-percentile value over an interval of ±15 s around each time point. We corrected the X-Y shift between imaging datasets with the NoRMCorre algorithm and then identified the same ROI in the different sessions with the register_ROIs function in the CaImAn package (https://github.com/flatironinstitute/CaImAn-MATLAB). Neurons that were identified as the same neuron in at least one early session and at least one late session were defined as pursued neurons.

We constructed a decoding model to predict the pole movement from the activity of a single neuron detected in both early and late sessions with the same regression model used for the one-photon imaging data. To estimate the change in cv*R*^2^ of the single-neuron decoding performance during learning, the cv*R*^2^ of each neuron was averaged over the early sessions or late sessions and the averaged cv*R*^2^ was compared between early and late sessions.

In addition to the single-neuron decoder, another decoding model was constructed to predict pole movement from the activities of multiple neurons in individual recording datasets. To reduce the possible influence of the difference in the number of trials and neurons between early and late sessions on the decoding performance, we randomly selected 10 pull/push trials and the same number of pursued neurons with early and/or late cv*R*^2^ of > 0.02 from early and late imaging sessions. The number of neurons used for the decoder construction was determined as the minimum number of pursued neurons in the early and late imaging sessions in the set. The numbers for individual datasets are shown in Fig. 6G, H. We excluded sessions with less than 10 pursued neurons from this analysis (imaging session 3 in marmoset 5 and session 1 in marmoset 6). To reduce overfitting by the decoder constructed from high-dimensional neuronal activity data, we then searched for the neuronal subspace that captured most of the variance in the original neuronal activity space as follows. First, the neural activity data during –1.0 to +0.13 s from the movement onset were trial-averaged and principal component analysis was used to generate a matrix that transformed the original neural activity to low-dimensional population activity capturing more than 95% of the variance in the trial-averaged activity. The low-dimensional activity in the individual trials was calculated by applying the same transformation matrix to the neural activity in the trials.

The predicted Y-axial trajectory at time *t*, *ŷ(t)* was expressed using the following formula:

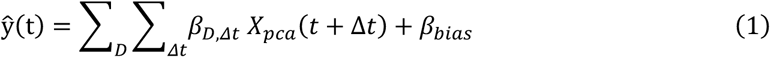

where *X_pca_* is the low-dimensional population activity, Δ*t* was set to 0, ± 0.1, ± 0.2, or ± 0.3 s, *βD*, Δ*t* is the coefficient for the low-dimensional activity in the *D*th dimension at Δ*t*, and *β*_*bias*_ is the bias term. *ŷ(t)* was calculated by 5-fold cross validation. To assess the changes in the cv*R*^2^ during learning, the cv*R*^2^ was averaged over early sessions or late sessions, and a Δpopulation decoder for cv*R*^2^ was calculated by subtracting the late sessions-averaged cv*R*^2^ from the early sessions-averaged cv*R*^2^. These processes started with a random selection of 10 pull/push trials and pursued neurons, and were repeated 1000 times, with the difference being considered statistically significant when the 2.5 percentile of the 1000 Δpopulation decoder cv*R*^2^ was above 0.0, or the 97.5 percentile was below 0.0.

Raw image sequences acquired with wide field-of-view two-photon microscopy were motion-corrected with the NoRMCorre package. A time-averaged image was used as the target image for the motion correction. Then, to improve the signal-to-noise ratio of the imaging data, we performed shot noise-reduction using the deep self-supervised learning-based denoising algorithm DeepCAD-RT (Li et al., 2021). All datasets were denoised with the convolutional neural network (CNN) that was trained on all datasets (11 imaging sequences from six imaging sessions). The training process was terminated at the 10th iteration because the performance of the trained CNN was optimal, as mentioned in the original article. The CNMF algorithm was employed to extract neuronal activities from a time-series of images (CaImAn package for MATLAB) (Pnevmatikakiset al., 2016). The factor of the autoregressive system was such that the inferred neuronal activities had one decay time constant in this step. We defined those ROIs whose automatically extracted activities reached the following criteria as the active neurons: a maximum spatial component size of 250; a minimum spatial component size of 10; a threshold for the spatial correlation between the estimated component and raw data of 0.6; a threshold for the temporal correlation between the estimated component and raw data of 0.5; a minimum signal-to-noise ratio of the estimated component of 2. For each active neuron, we calculated the detrended relative fluorescent change (Δ*F*/*F*) using the detrend_df_f function in the CNMF package with a time window of ±15 s for calculating background fluorescence. The single-neuron decoder was constructed with the same protocol as described above, but the pole trajectory was down-sampled to 7.5 Hz and Δ*t* was set to 0, ± 0.133, ± 0.266, or ± 0.399 s, because the frame rate of the wide field-of-view two-photon imaging was 7.5 Hz.

### Tracking of body movements

Movements of the upper limbs, tongue, and pupil in the videos were tracked with DeepLabCut (Mathis et al., 2018). Fifty frames from video representing the dataset were automatically extracted on two experimental days. Since the tongue was not visible in many frames for marmosets 1 and 2, another 50-frame dataset that included the tongue movement was manually prepared from the same video. In the extracted dataset, the position of each part of the body was manually labeled, and on the basis of this labeled dataset, the position of each body part in the other frames was predicted with DeepLabCut. We adopted the data with a likelihood of > 0.95 (for the marmosets 1 and 2) or > 0.6 (for the marmosets 3–6), and the movement in the discarded frames was linearly interpolated using the data before and after these frames. The position of the pupil was determined as the midpoint of the predicted positions of the left and right edges of the pupil. Pupil diameter was calculated as the distance between these positions. The predicted position of each part of the body was smoothed with a moving average of five frames. The trajectories of the pole, left hand, right hand, left elbow, left shoulder, left knee, right knee position, pupil position, and pupil diameter were z-scored in the same way as Δ*F*/*F*. As described above, the images for marmosets 1 and 2 had a lower pixel number and a higher frame rate (shorter exposure time) than the images for marmosets 3–6. Thus, the quality of the former images was worse than that of the latter images, meaning that the tongue position when the mouth was closed was frequently assigned to the wrong place in marmosets 1 and 2. Therefore, the following criteria to detect licking were also set for marmosets 1 and 2: the predicted tongue position should be inside the ROI set near the mouth, and the intensity of another ROI over the mouth should be above a threshold value (for marmoset 1, the 30th percentile of the intensity values during the imaging; for marmoset 2, the 50th percentile of the values) because the tongue showed higher intensity than the closed mouth. No licking was assumed during periods when these criteria were not met.

### Statistical analyses

Statistical analyses were performed using MATLAB (2018–2020a, MathWorks). The Wilcoxon signed-rank test, Wilcoxon rank sum test, Spearman’s rank correlation coefficient test, Pearson’s rank correlation test, and random permutation tests were used for statistical comparisons. No statistical tests were run to predetermine the sample size. Data are presented as mean ± SEM unless otherwise noted.

### Data availability

All data and computer codes are available from the corresponding author.

