## Supplementary Information for "Dynamics of motor direction representation in the primate premotor and primary motor cortices during sensorimotor learning"

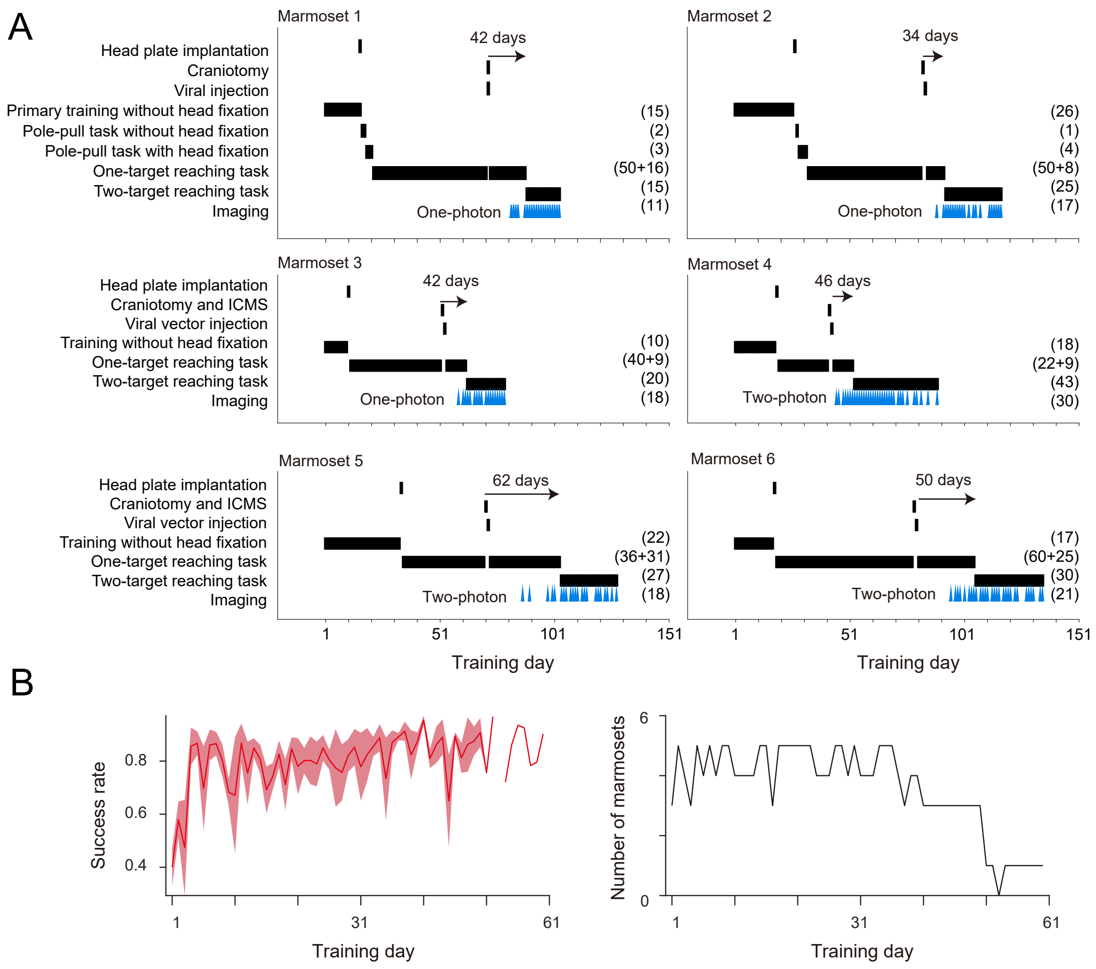


**Figure S1. The task schedules for the six marmosets.**
(A) Black bars indicate the experimental days for the surgeries and tasks that are indicated on the left for each marmoset. The numbers in parentheses on the right indicate the total session numbers for the corresponding task or imaging. The days above the arrows indicate the interval between the virus injection and the first session of the TTR, which includes non-training days. Blue arrowheads represent the sessions with imaging experiments (imaging sessions).
(B) Time course of the rate of successful trials to total pull trials in the OTR task sessions. The right image shows the number of animals in each training session. Since some of the data are missing, the number sometimes increases or decreases across sessions. The shading indicates ± SEM.
The format corresponds to that of Figure 1D.

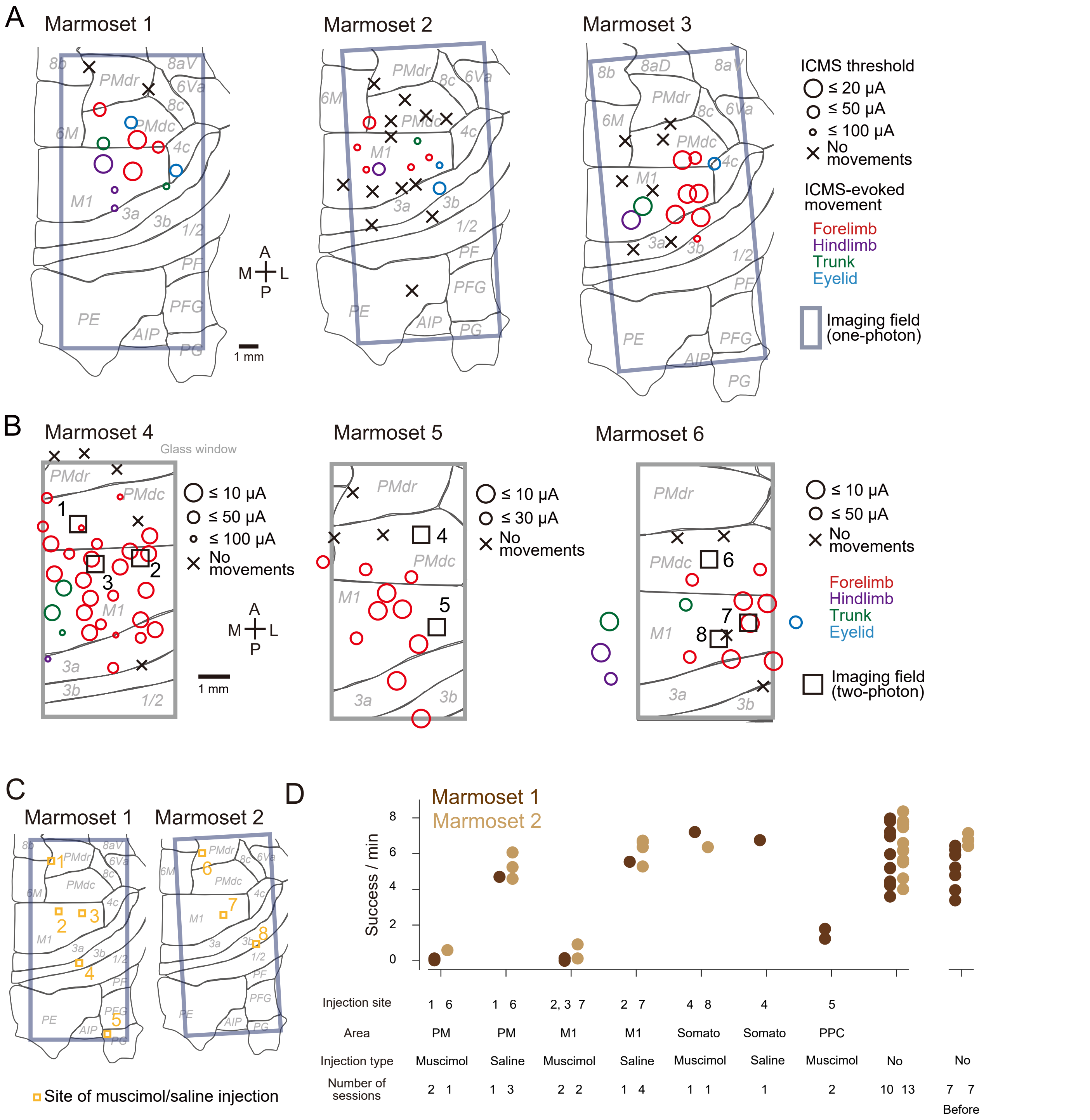


**Figure S2. Results of the ICMS and muscimol injection experiment.**

(A, B) Motor mapping by ICMS in marmosets 1–3, which were used for chronic one-photon imaging experiments (A), and in marmosets 4–6, which were used for chronic two-photon imaging experiments (B). In (B), the black boxes indicate the imaging fields of the two-photon imaging area. Movements of forelimb (red), hindlimb (purple), trunk (green), and eyelid (cyan) were identified by visual inspection. The circle diameter indicates the amplitude of the current needed to induce the movement. A, anterior; P, posterior; M, medial; L, lateral.

(C) Muscimol injection sites (yellow boxes) in marmosets 1 and 2.

(D) The number of successful trials per minute in the TTR task in each condition of the injections in marmosets 1 (dark brown) and 2 (light brown). PM, premotor cortex (sites indicated by 1 and 6 in C); M1, primary motor cortex (sites indicated by 2, 3, and 7 in C); PPC, posterior parietal cortex (site indicated by 5 in C); Somato, somatosensory cortex (sites indicated by 4 and 8 in C). The performance in the sessions without the injection is indicated in the No injection columns. The two right-most columns indicate the number of successful trials per minute in seven training sessions just prior to the injection experiment. The results of the sessions 1 day after the injection and sessions when the training stopped within 5 minutes are excluded from this analysis.

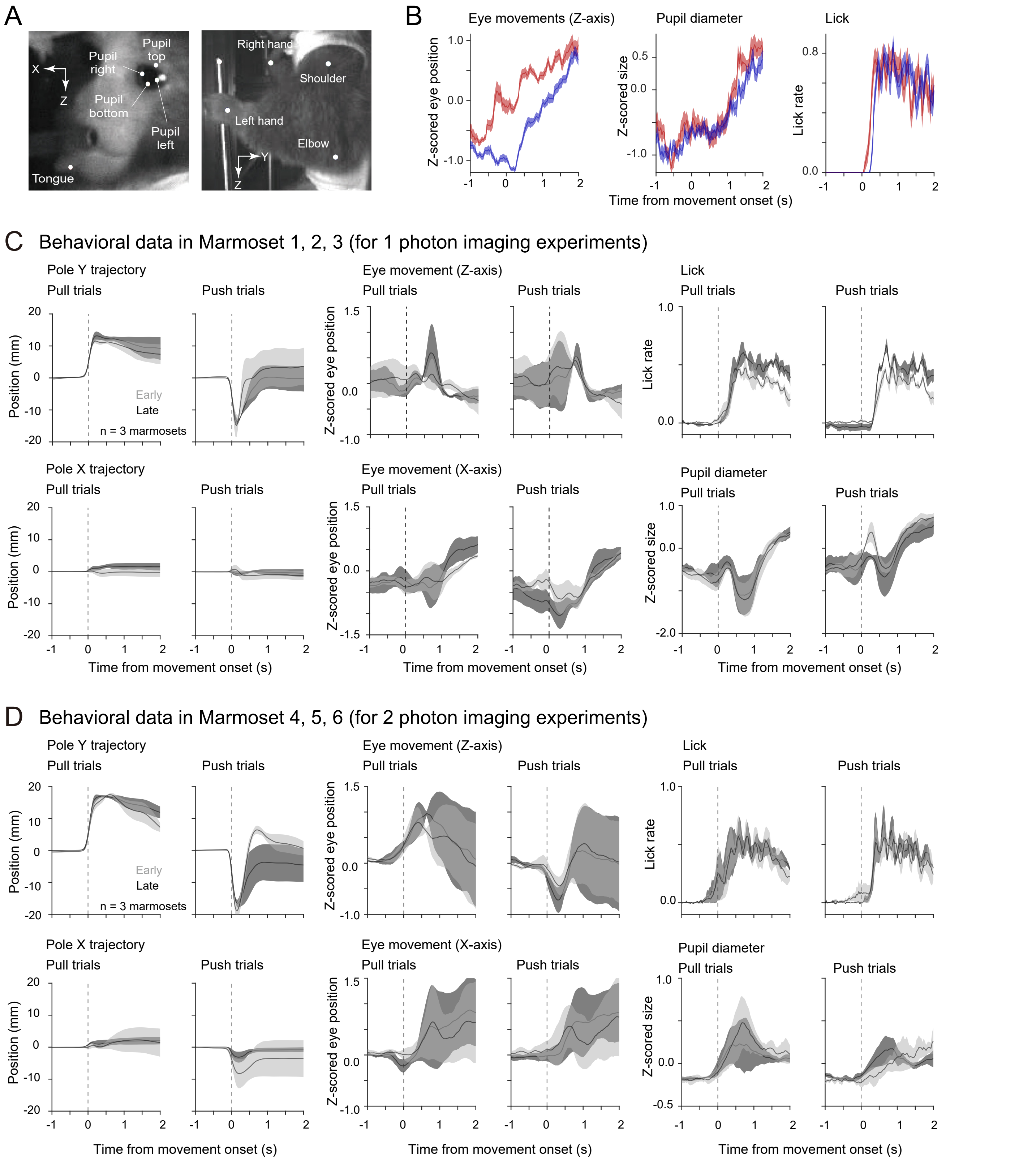


**Figure S3. Behavioral changes from early to late sessions.**

(A) Representative images of the face and upper body from the two cameras. The points of the extracted body parts are overlaid.

(B) Representative trial-averaged time course of the eye position along the Z-axis (vertical axis), pupil diameter, and lick rate. The traces are aligned to the onset of the pull/push pole movement.

(C, D) Trial-averaged time courses of the Y-axial pole position, Z-axial eye movement, lick rate, X-axial (right-left axis) pole trajectory, X-axial (right-left axis) eye movement, and pupil diameter for successful pull and push trials in the early (light gray) and late (dark gray) sessions for marmosets 1–3 (B) and 4–6 (C). All traces are aligned to the onset of the pole (pull or push) movement. Shading indicates ± SEM (n = 3 animals).

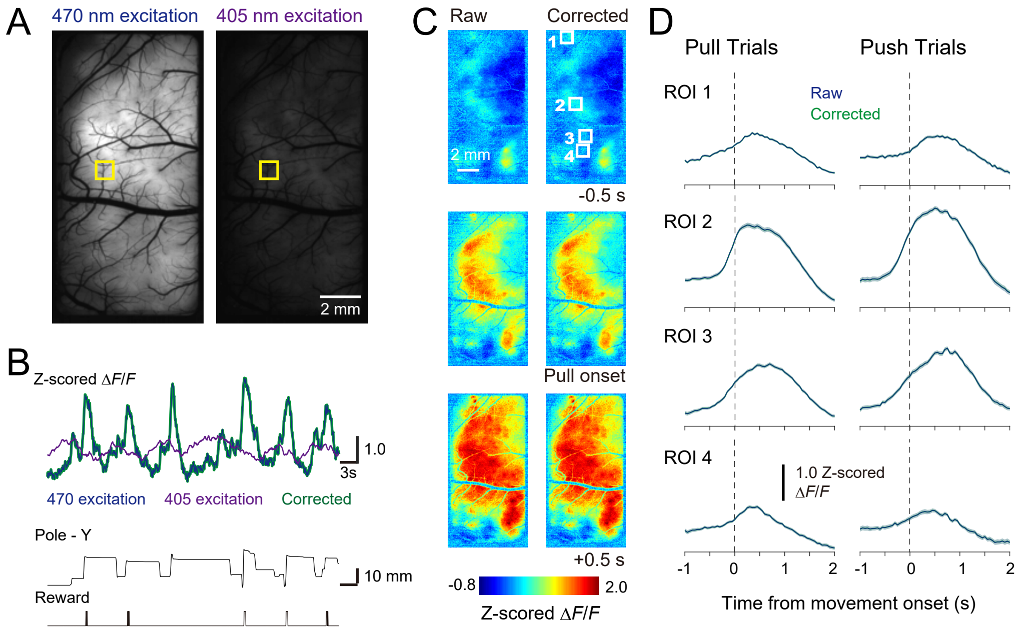


**Figure S4. Hemodynamic signals were negligible in one-photon calcium imaging.**
(A) Images of GCaMP6s fluorescence with 470 nm (left) and 405 nm (right) light excitation.
(B) Neuronal and hemodynamic responses during the task performance. Blue line indicates z-scored blue light-excited $\Delta F/F$ of the pixels inside the ROI (yellow box in A). Violet line indicates the z-scored smoothed violet-light-excited $\Delta F/F$ of the same ROI. Green line represents the hemodynamic corrected $\Delta F/F$. The Y-axial pole trajectory and the reward timing during the task performance are indicated in the bottom image. Note that the blue and green lines almost overlap.
(C) Trial-averaged blue light-excited $\Delta F/F$ images and the corrected z-scored $\Delta F/F$ images at –0.5 s, 0.0 s, and +0.5 s from the pull onset.
(D) Trial-averaged blue light-excited $\Delta F/F$ traces and corrected $\Delta F/F$ traces aligned to the pull and push onsets. Blue and green lines indicate the blue light-exited and corrected $\Delta F/F$ of ROIs, respectively, as indicated in C. Note that the blue and green lines almost overlap.

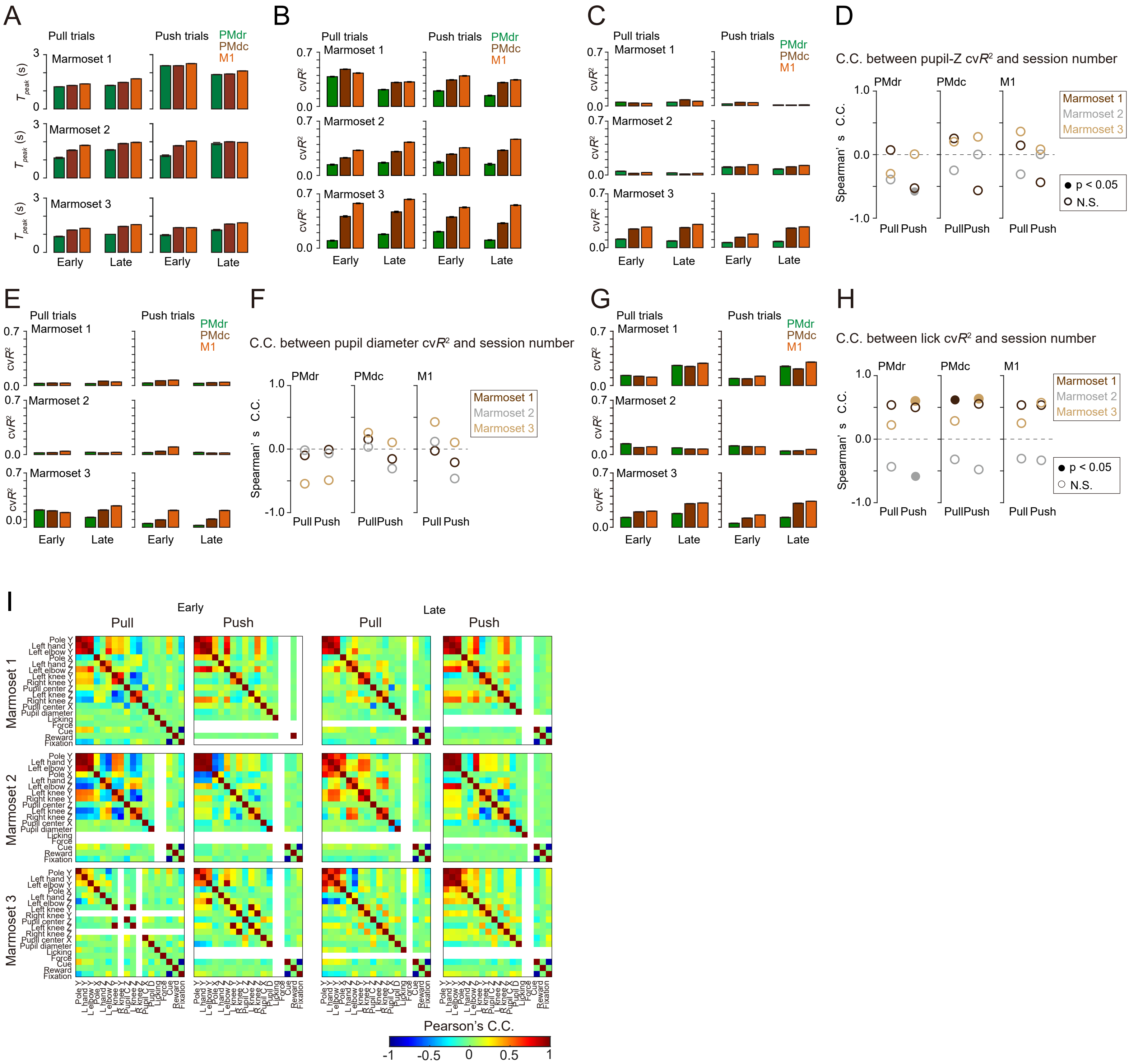


**Figure S5. $T_{peak}$ and prediction accuracy of the orofacial movements and correlation matrices between behavioral variables.**
(A) $T_{peak}$ of PMdr (green), PMdc (brown), and M1 (orange) for pull and push trials in the early and late sessions of marmosets 1–3.
(B, C, E, G) cv$R^2$ for the pole-Y trajectory (B), Z-axial eye movement (C), pupil diameter (E), and lick rate (G) of PMdr, PMdc, and M1 for pull and push trials in the early and late sessions of marmosets 1–3.
(D, F, H) Spearman' s rank correlation coefficients between cv$R^2$ and Z-axial eye movement (D), pupil diameter (F), and lick rate (H), and according to imaging sessions, in marmosets 1–3. Closed circles, $p < 0.05$ for the correlation coefficients. Open circles, $p > 0.05$. The numbers of imaging sessions were 11, 17, and 12 for marmosets 1, 2, and 3, respectively.
(I) Correlation matrices between the behavioral variables for pull and push trials in the early and late sessions of marmosets 1–3. Pearson' s correlation coefficients between the behavior data during –1.0 to 0.13 s from the movement onset were calculated for each session and averaged across early or late sessions. We excluded the correlation coefficients between two variables when one or both of the variables did not show any change during the time window in all early or late sessions (white).

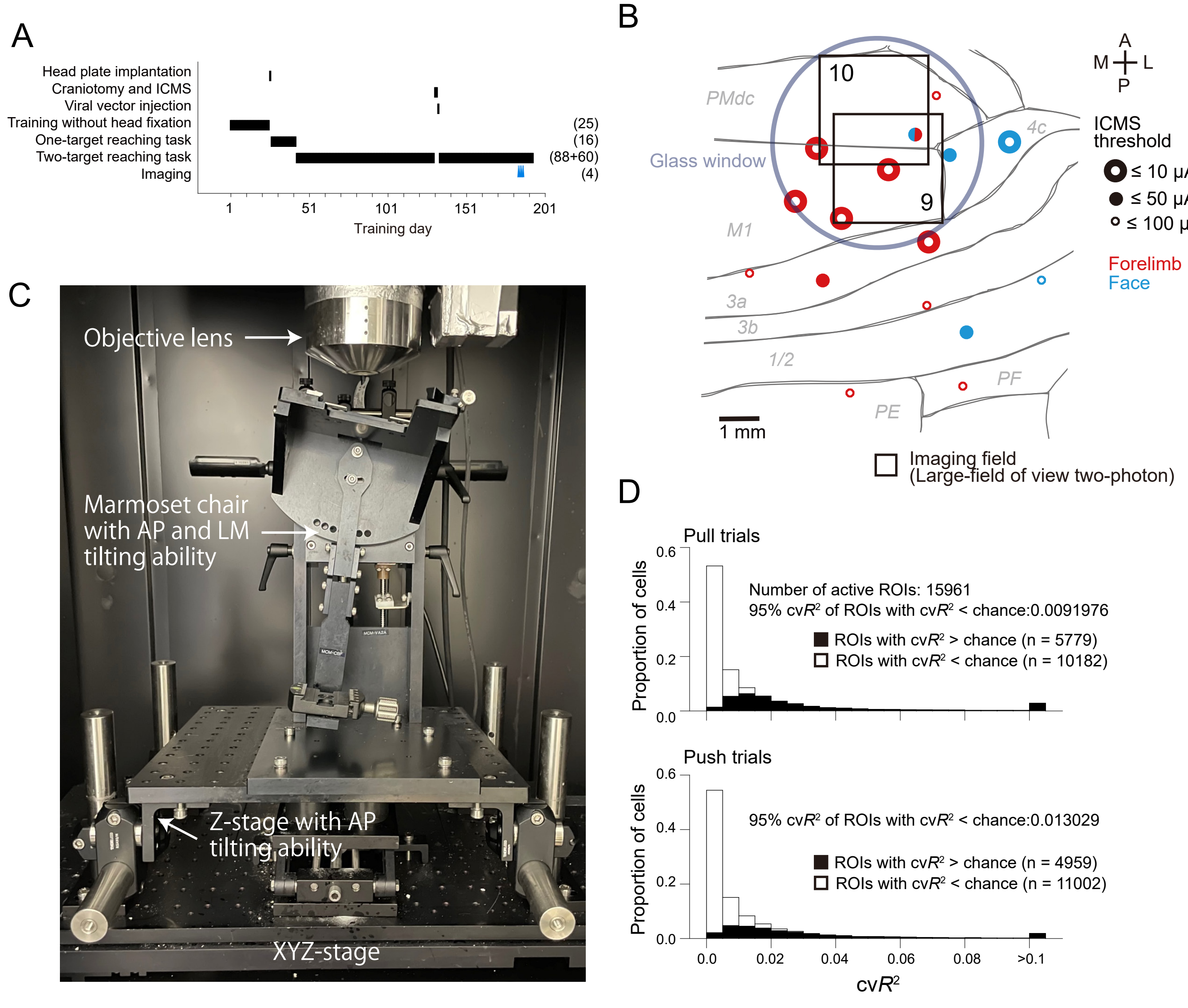


**Figure S6. The experimental and analytical data related to the wide field-of-view two-photon imaging.**
(A) The task schedule for marmoset 7. The conventions are the same as in Figure S1A.
(B) Results of ICMS in marmoset 7 and the location of the glass imaging window. Inferred borders of the cortical areas are overlaid. The imaging fields are shown as two boxes (see also Table S1). The conventions are the same as in Figure S2B.
(C) The marmoset chair and stage under the objective lens for wide field-of-view two-photon calcium imaging. To set the imaging field perpendicular to the objective lens, the marmoset chair was tilted 24° along the lateral-to-medial axis and 16° along the anterior-to-posterior axis for imaging in marmoset 7.
(D) Histogram of the cv$R^2$ of individual active neurons for successful pull (top) and push (bottom) trials. The data were pooled from the six imaging sessions. Closed bars indicate the neurons whose cv$R^2$ was higher than the top 95th percentile value of cv$R^2$ calculated from the trial-shuffling data. Open bars indicate the other neurons.

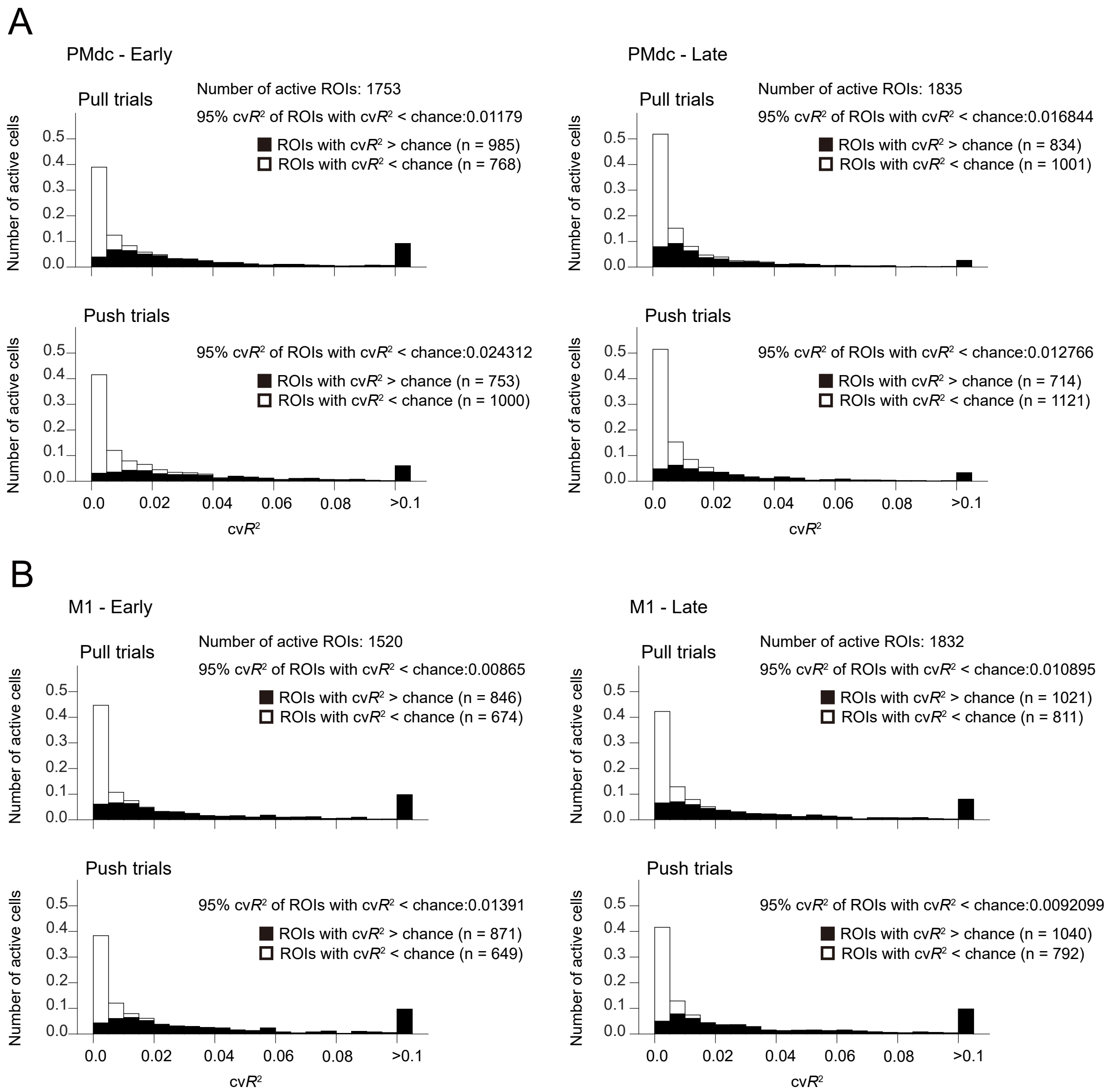


**Figure S7. Distribution of and changes in the cvR2 of individual neurons.**
(A, B) Histogram of the cv$R^2$ of individual active neurons for successful pull and push movements in early and late sessions from three PMdc areas (A) and three M1 areas (B) from marmosets 4–6 (total of 34 imaging sessions). The conventions are the same as in Figure S6D. In Fig. S6D, S7A, and S7B, almost all neurons with cv$R^2$ of > 0.02 had higher cv$R^2$ values than the top 95th percentile value of cv$R^2$ calculated from the trial-shuffling data. Therefore, we defined these neurons as pull- and/or push-related neurons.

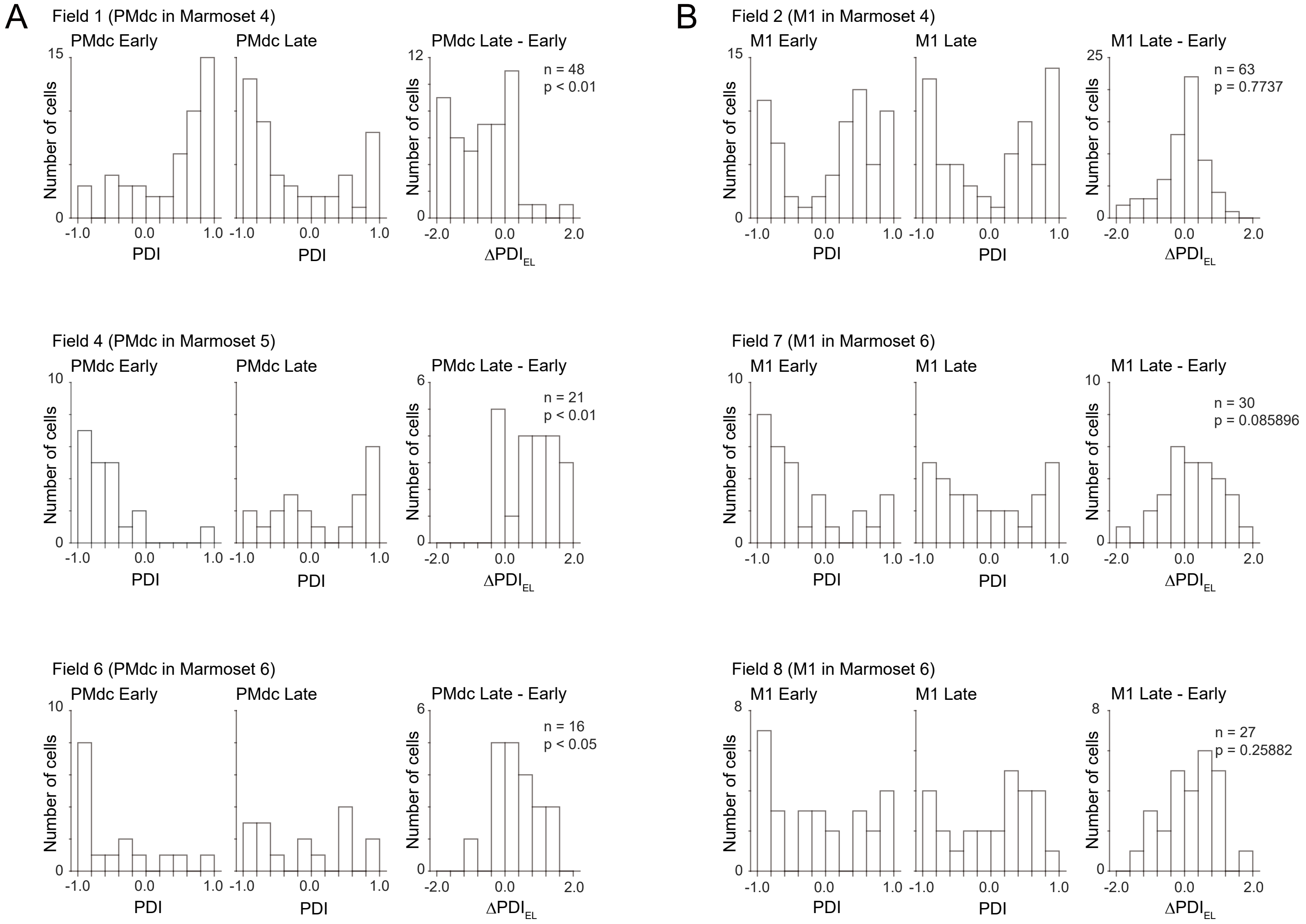


**Figure S8. Histograms of the PDI of neurons and the difference in PDI in each imaging field.**

(A, B) Histograms of PDI of the pursued neurons in early (left) and late (middle) sessions and $\Delta PDI_{EL}$ (right) in each imaging field of PMdc (A) and M1 (B). P-values are shown for Wilcoxon signed-rank tests comparing the PDI of the neurons between early and late sessions. Only pursued neurons that were pull-related and/or push-related in both early and late sessions were included in this analysis (the same as in Fig. 7A and B).

**Table S1. List of the training sessions for calcium imaging experiments.**

| Session | Recording method | Imaging field (cortical depth) |
|---|---|---|
| Marmoset 1 | | |
| 1, 2, 3, 4, 5, 6, 7, 8, 9, 10, 11, 12, 13, 14, 15 | 1P | |
| Marmoset 2 | | |
| 1, 2, 3, 4, 5, 6, 7, 8, 9, 11, 13, 14, 16, 20, 21, 22, 23, 24, 25 | 1P | |
| Marmoset 3 | | |
| 1, 2, 4, 5, 6, 7, 9, 10, 11, 12, 13, 14, 15, 16, 17 | 1P | |
| 18, 19, 20 | 1P, 1P hemodynamic | |
| Marmoset 4 | | |
| 1, 3, 5, 7, 9, 11, 13, 15, 17, 20, 22, 27, 30, 33 | 2P in PM | 1 (250 μm) |
| 6, 8, 10, 12, 14, 16, 18, 21, 24, 28, 37 | 2P in M1 | 2 (250 μm) |
| 2, 4, 6 | 2P in another M1 | 3 (250 μm) |
| Marmoset 5 | | |
| 1, 3, 7, 10, 12, 17, 20, 21, 23, 25 | 2P in PM | 4 (215 μm) |
| 2, 5, 6, 8, 11, 16, 18 | 2P in M1 | 5 (215 μm) |
| Marmoset 6 | | |
| 5, 6, 8, 10, 11, 13, 14, 23, 25, 30 | 2P in PM | 6 (275 μm) |
| 1, 4, 6, 9, 11, 14, 16, 19, 24, 26, 30 | 2P in M1 | 7 (275 μm) |
| 3, 4, 5, 8, 10, 13, 15, 23, 24, 29 | 2P in M1 | 8 (240 μm) |
| Marmoset 7 | | |
| 1, 3 | 2P in M1 and PM | 9 (150 and 250 μm) |
| 2, 4 | 2P in M1 and PM | 10 (150 and 250 μm) |

1P, one-photon imaging; 2P, two-photon imaging; PM, premotor cortex; M1, primary motor cortex. The number in the imaging field column corresponds to that in Fig. S2A, S2B, and S6B. Session numbers with the wavy and solid underlines indicate the early and late sessions, respectively. Session numbers with the gray hatching indicate the sessions that were removed from the analysis because these sessions included less than 10 pull or push trials. The sessions indicated with gray letters were not analyzed because of the following reasons: In sessions 30 and 33 of the marmoset 4, the fluorescence intensity of ROIs was apparently lower than those in other sessions. In session 6 of the marmoset 4, the size of the FOV in the site 2 was different from that in the other sessions. In the imaging field 3 of the marmoset 4 and imaging field 5 of the marmoset 5, the imaging was not conducted at the late stage of the training sessions. In each of the field 9 and 10 in marmoset 7, the imaging was conducted in 2 sessions in the same field at the similar cortical depth. We excluded the two sessions of the imaging field 10 at the depth of 250 mm from the analysis because the number of active and movement-related ROIs in these datasets were apparently lower than those in other sessions: 18.4 and 20.3 movement-related ROIs per $mm^2$ were detected in these datasets, while 51.7–158.9 movement-related ROIs per $mm^2$ in other datasets in marmoset 7 and 69.4–344.7 movement-related ROIs/$mm^2$ in chronic imaging datasets. We used all other six datasets in marmoset 7 for the analysis in Fig. 5 and S6 because not the same population of neurons were recorded in the corresponding sets of the experiments due to slight changes in the tilting angle of the marmoset chair.
